# Mitochondrial DNA heteroplasmy drives cortical neuronal disturbances in human organoids harbouring the common m.3243A>G mutation

**DOI:** 10.1101/2025.03.21.644499

**Authors:** Denisa Hathazi, Camilla Lyons, Daniel Lagos, Oliver Podmanicky, Mariana Zarate-Mendez, Nell Nie, Juliane S Müller, Kieren S J Allison, Majlinda Lako, Eva Morava-Kozicz, Tamas Kozicz, Patrick Chinnery, András Lakatos, Rita Horvath

## Abstract

Mitochondrial diseases frequently affect the brain leading to severe and disabling neurological symptoms. The heteroplasmic m.3243A>G mutation in *MT-TL1*, encoding mt-tRNA^Leu^, is responsible for ∼80% of mitochondrial encephalomyopathy, lactic acidosis, and stroke-like episodes (MELAS), which is one of the most characteristic mitochondrial syndromes, leading to disability and early death. There are no animal models harbouring this mutation to provide precise mechanistic insights informing therapeutic interventions. Here, we generated a human iPSC-derived cerebral organoid slice model that recapitulates cortical architecture and mitochondrial pathology. Using biological assays and single-cell RNA sequencing, we uncovered heteroplasmy-dependent transcriptional shifts and changes in key cellular processes in cortical neurons. Organoids with high heteroplasmy showed a predominant impairment of deep-layer neurons triggered by mitochondrial stress, leading to axonal degeneration and apoptosis, similar to brain autopsy of a MELAS patient. Our findings provide insights into the vulnerability of long-range projection neurons in mitochondrial diseases, advancing our understanding of disease mechanisms with a view to potential therapeutic strategies.

## INTRODUCTION

Mitochondrial diseases are clinically heterogeneous genetic conditions, caused by the defect of energy production through oxidative phosphorylation (OXPHOS)^1^. Neurological manifestations are common and severely disabling, given that neuronal function relies on ATP generated through OXPHOS^2^. The mechanisms by which mitochondrial mutations lead to neuronal defects and disease phenotypes are poorly understood, impeding the development of therapeutic strategies. Discoveries have been hampered by the complexity of human mitochondrial genetics and the lack of reliable animal and cellular models.

Studying the downstream effects of mitochondrial DNA (mtDNA) mutations has been particularly challenging due to the variable heteroplasmy levels that characterize human mitochondrial diseases, such as Mitochondrial Encephalomyopathy with Lactic Acidosis and Stroke-like Episodes (MELAS) ^3,4^. While mouse models have been successfully developed to study nuclear-encoded mitochondrial disorders^5^, generating heteroplasmic mtDNA mutations in mice has proven difficult with the few existing mtDNA mouse models failing to recapitulate the neurological pathologies observed in patients^3,4^. Recent advances in single-nucleotide mtDNA editing have enabled the generation of heteroplasmic variants in cellular and animal models^5–8^. However, these techniques do not allow for transgenic manipulation of all mtDNA mutations. Importantly, there are no animal models available for the heteroplasmic m.3243A>G variant, one of the most common mtDNA variants^9^.

In fact, the m.3243A>G mutation in the *MT-TL1* gene has been detected in almost half of adult patients with mitochondrial disease in clinical cohorts presenting with broad phenotypes of maternally inherited diabetes and deafness (MIDD), chronic progressive ophthalmoparesis (CPEO) and mitochondrial myopathy, often accompanied by disabling neurological symptoms, including ataxia, peripheral neuropathy and Stroke-like Episodes (SLEs)^10^. Thus, there is an unmet need for human-based research platforms to study its effects and develop targeted therapies.

Recent *in vitro* technologies enabled studies using human neuronal models harbouring the m.3243A>G mutation^11^. A few studies involved human induced pluripotent stem cell (hiPSCs)-derived glutaminergic neurons with m.3243A>G, including iNeurons^12,13^, motor neuron (MN) progenitors and MNs^14^, describing oxidative phosphorylation induced structural and metabolic disturbances. Distinct changes had been also described for other cell types, such as fibroblasts and hiPSCs carrying m.3243A>G^11,13^. Therefore, how the varying degree of heteroplasmic mtDNA affect various somatic neural cells in a tissue environment is a pertinent question to address, which may shed light on key targetable downstream mechanisms.

Three-dimensional (3D) hiPSC-derived multicellular neural organoid models offer new promise for cell type-specific mechanistic discoveries in human neurological disease and treatment strategies^15,16^. Cortical organoid slice culture systems allow long-term assessments, perturbations and easy access for monitoring outcome measures in several neural cell types^17^. In addition, 3D organoids allow neurons to develop within a structured tissue-like environment, preserving key cell–cell interactions and spatial gradients essential for neuronal diversification and maturation^18,19^. Specifically, the presence of layer specific cortical neuronal identities in organoids, including deep- and upper-layer subtypes, is an advantage over 2D monolayer culture models in which simultaneous divergence of neuronal subtypes seldom occurs^17,19^.

Human cortical organoid models have not yet been generated to uncover the role of m.3243A>G heteroplasmy in orchestrating cellular pathology in various neuronal subtypes. To overcome this problem, we generated 756, 3D cortical organoid slice cultures from 7 iPSC lines, derived from two patients with the m.3243A>G variant within 5 batches. We used a combination of single-cell transcriptomic approaches and cell cultures-based assays to reveal heteroplasmy-driven cellular changes, with a focus on neuronal subpopulations. We show that organoid slices display consistent tissue composition and stable heteroplasmy rates associated with mitochondrial dysfunction. Specifically, we provide direct evidence that high m.3243A>G heteroplasmy leads to impaired bioenergetics, disturbed neuronal structure and function, associated with a significant loss of cortical deep layer neuronal population in organoids, similar to postmortem human adult brain of patients with MELAS.

## RESULTS

### m.3243A>G hiPSC-derived organoids have homogeneous cell populations and retain the mtDNA mutation

To assess the fidelity of human cortical organoids to recapitulate neural tissue architecture and the diversity of cell types harbouring the heteroplasmic m.3243A>G variant, we employed a previously characterized long-term organoid slice culture platform^17,19^. For this, we used iPSC lines derived from two patients with the m.3243A>G mutation (Figure 1A), which were subjected to clonal isolation and expansion based on their heteroplasmy levels. For patient 1, we developed three clones with heteroplasmy levels of 12%, 58%, and 72%. For patient 2, we generated four iPSC clones with 0%, 15%, 42%, and 64% heteroplasmy (Figure S1 A). The clones with the lowest heteroplasmy - 12% for patient 1 and 0% for patient 2 - were below the mutational threshold and were used as isogenic controls. When assessed by phase contrast microscopy embryonic bud formation was comparable between all heteroplasmic organoids and their isogenic controls at 10 days in vitro (DIV) (Figure S1 B). We performed immunohistochemical analyses to verify potential heteroplasmy variation-driven differences in tissue architecture in 30 DIV organoids. We observed the germinal zones in all heteroplasmy groups, with the presence of the typical early neuronal cell phenotypes, including SOX2+ neural epithelium, NEST+ progenitors, and TUJ+ immature neurons (Figure 1B and Figure S1C). This consistent tissue organisation allowed us to set up organoid slice cultures, which can be cultivated long-term to quantitatively compare heteroplasmy-induced effects imposed on mature neurons. Organoids were sliced between 45-50 DIV and grown further until 100 DIV as air-liquid interface cortical organoid (ALI-CO) slices, as previously described^17,19^, which allows consistent cortical expansion, and neuronal maturation. Normal development in mutant organoids mirrored normal brain development and later symptom onset seen in patients with m.3243A>G.

**Figure 1.**
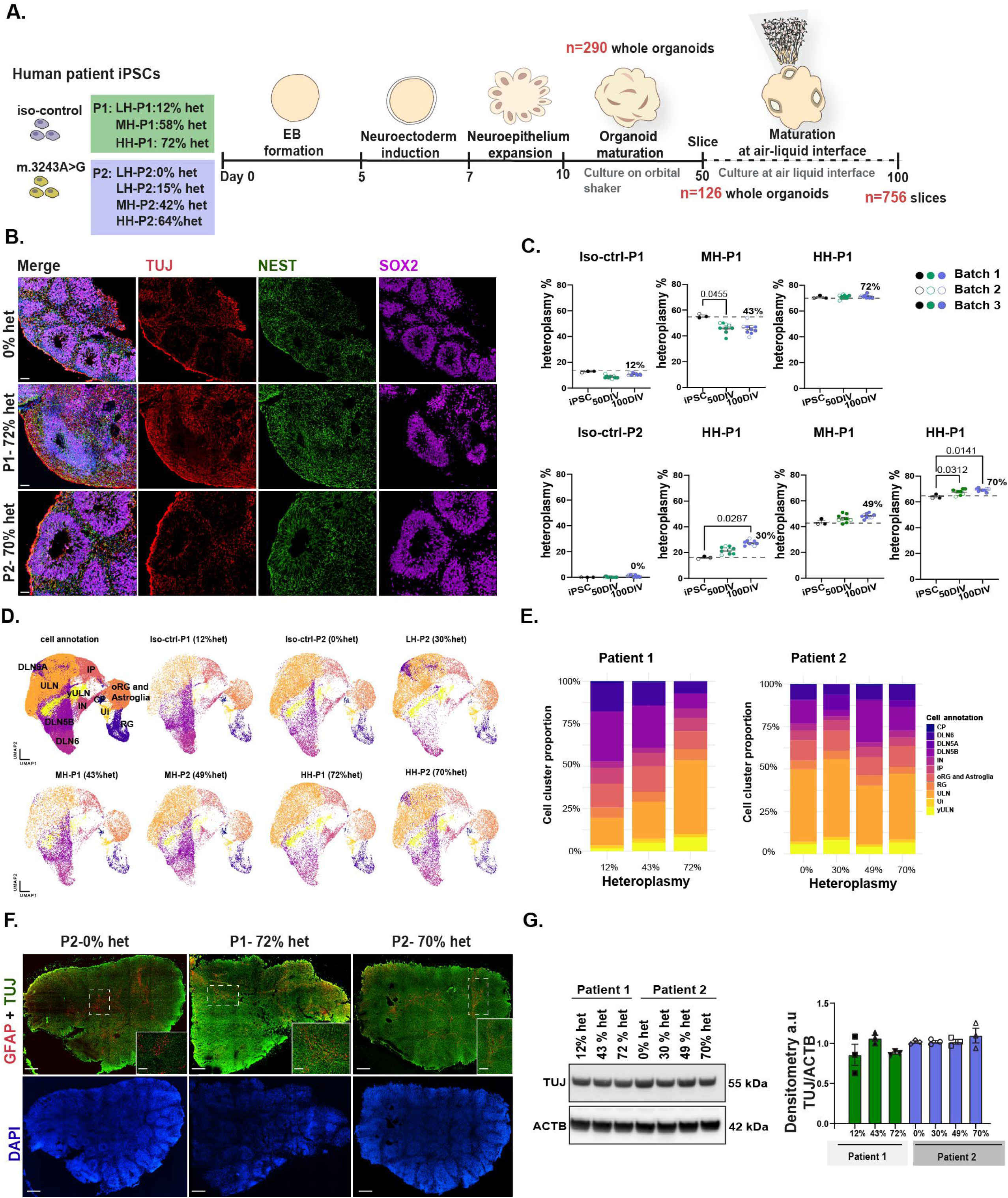
m.3243A>G ALI-COs retain the mutation and exhibit consistent cellular populations. **A.** Schematic representation of the organoid generation process and the human iPSC lines used in this study; **B.** Representative immunofluorescence images of cortical organoids at DIV 30, showing the palisade-like organization of SOX2LJ progenitor cells (violet), surrounded by immature TUJLJ neurons (red) and NESTINLJ progenitor cells (green). Scale bar = 50 µm; **C.** Heteroplasmy levels were assessed in the starting iPSCs and cortical organoids at DIV 50 and 100 using pyrosequencing. For organoids, heteroplasmy was measured across three independent batches, with three organoids analysed per batch to account for intra- and inter-batch variability *n* = 3 for iPSCs and *n* = 9 for organoids. Data represent the mean ± SEM; Statistical analysis: one-way ANOVA, mixed-effects analysis with Dunnett’s post hoc test (overall ANOVA p-values indicated in the graphs); **D.** Uniform manifold approximation and projection (UMAP) plots illustrating 11 color-coded clusters identified in the scRNA-seq analysis, presented for the merged dataset and for each individual heteroplasmic organoid (nLJ=LJ119,591 cells). Clusters were defined using cell-marker genes and include the following populations: ULN, upper-layer cortical neurons; DLN5B and DLN5A, deep-layer neurons (layer 5, subtypes A and B); DLN6, deep-layer neurons (layer 6); IN, interneurons; yULN, young upper-layer neurons; IP, intermediate progenitors; RG, radial glia; oRG, outer radial glia; CP, choroid plexus; Ui, unidentified cells; **E.** Bar plots depicting the cluster identity distributions, expressed as cell proportions within each pooled patient pair sample; **F.** Representative immunofluorescence images of ALI-COs at DIV 100, showing TUJLJ (green) neurons and GFAPLJ (red) glia across three biological replicates per group. Scale bar = 200 µm; **G.** Representative WB images (left) of TUJ and ACTB protein expression in ALI-COs at DIV 100, along with quantification (right) of TUJ band intensity normalized to ACTB. Data represent the mean ± SEM; *n* = 3 independent ALI-COs per heteroplasmic line and isogenic controls from different batches. Statistical analysis: one-way ANOVA with Tukey’s post hoc test (overall ANOVA p-values indicated in the graphs).

We then explored the cell type composition and maturity to uncover any biases that may affect the assessment of heteroplasmy levels and metabolic function across the different groups of organoids. Since heteroplasmy can change over time, we quantified these levels at 50 DIV (post-slicing) and at 100 DIV using pyrosequencing, comparing the results to the initial mutation load detected in iPSCs (Figure 1C). While the absolute heteroplasmy levels slightly changed over time in patient organoids, each clone remained within its originally assigned range (i.e., low, medium, or high). In Patient 1, heteroplasmy decreased slightly from 58% to 42% by 100 DIV in one clone, while the high heteroplasmy clone maintained a steady level of 72%. For Patient 2, the low heteroplasmy organoids increased from 15% to 30% by 100 DIV, whereas the medium (from 42% to 49%) and high (from 64% to 69%) heteroplasmy clones showed only a minor increase.

The consistency in heteroplasmy ranges allowed us to effectively assess the impact of the m.3243A>G mutation load on cell type diversity and homogeneity using single-cell RNA sequencing of 100 DIV organoids. Clustering was performed on non-batch corrected merged samples using Seurat (Figure 1D), which identified 11 clusters corresponding with major cell types based on known marker genes apart from one unidentified cluster (Figure 1D, E).

All identified cellular populations were present in both patient-derived organoids, with similar relative proportions observed across individual samples and patients (Figure 1D, E). Moreover, immunolabeling analyses showed that the distribution of neurons and the levels of the TUJ neuronal marker were consistent among organoids with varying heteroplasmy (Figure 1F, G; Figure S1D). Together, these results indicate that heteroplasmy does not significantly affect cellular composition, as both the overall and neuronal cell proportions remain constant within the ranges of mtDNA mutation load.

### Neuronal diversity and maturity are not influenced by heteroplasmy levels

We investigated whether transcriptomic signatures distinguish distinct cellular subtypes and whether characteristics of these subtypes are affected by heteroplasmy. By 75 days in vitro (DIV), the telencephalic marker FOXG1 showed widespread immunoreactivity in organoids, and by 100 DIV it is broadly expressed across all cellular clusters, confirming forebrain identity (Figure 2A,B; Figure S2A). Corresponding with the early developmental stage at 100 DIV, immature cells, such as outer radial glia and astroglial cells (oRG/astroglia, marked by HOPX, TNC, GFAP or AQP4), inner radial glia (iRG, marked by TOP2A), and intermediate progenitors (IP, marked by EOMES) have been detected (Figure S2B and C). Our analysis identified signatures of multiple immature and mature neuronal cell types including excitatory upper layer neurons (ULN), marked by CUX2, SATB2, and DCX expression (Figure 2C and Figure S2B and C). Deep layer neurons (DLN) with makers for layer 5a, 5b and 6 were also identified based on the expression of FEZF2/CTIP2, CTIP2/PCP4 and TLE4, FOXP2, respectively (Figure 2C and Figure S2B). In addition, cells with interneuron (IN) identities characterised by GAD2 and DLX2 expression were also present (Figure 2SB). The cell-subtype-specific transcriptomic signatures suggest that overall cell identities remain largely unchanged, regardless of heteroplasmy levels.

**Figure 2.**
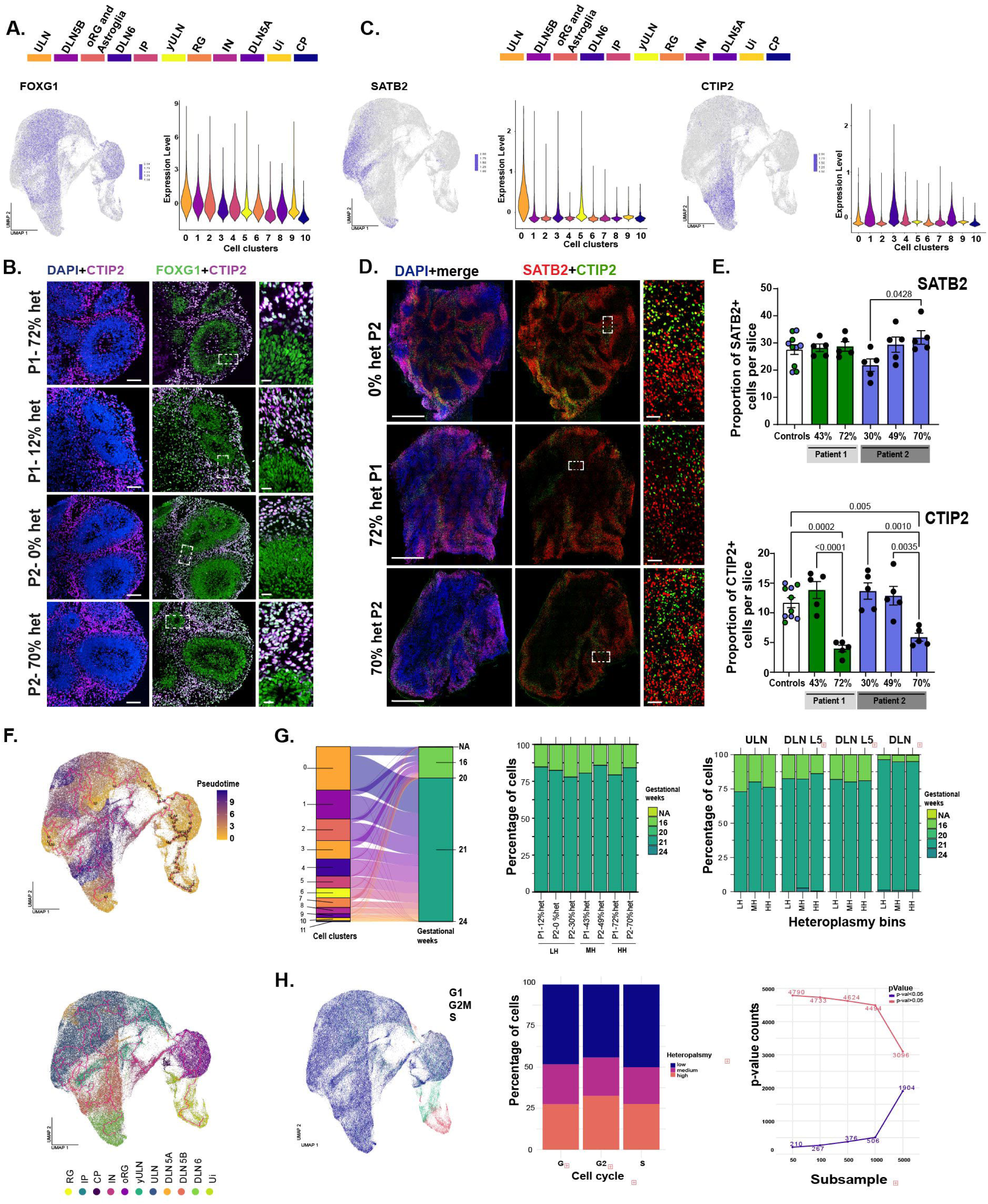
Cortical cell diversity and maturation are unaffected by heteroplasmy and are consistently recapitulated across all heteroplasmic organoids. A. and. **C.** Violin plots (left) and UMAP plots (right) illustrating the expression of FOXG1 (forebrain identity), CTIP2 (deep-layer neurons, DLN), and SATB2 (upper-layer neurons, ULN) marker genes identified by scRNA-seq in DIV 100 organoids. Cluster numbers (x-axis) correspond to color-coded cell type identities as indicated in the UMAP plots; **B.** Representative immunofluorescence images of cortical plate regions at DIV 75, showing the forebrain identity marker FOXG1 (green) and early-born CTIP2LJ neurons (violet), with insets highlighting cortical plate organization. Scale bar = 50 µm; **D.** Representative immunofluorescence images of laminar organization in cortical organoid slices at DIV 100, where SATB2LJ cells (red) mark ULNs and CTIP2LJ cells (green) label DLN5 neurons with scale bar= 200 µm; **E.** Bar plots quantifying SATB2LJ and CTIP2LJ cells, normalized to the total number of cells across the entire ALI-CO cryosection. Data represent the mean ± SEM; *n* = 5 independent ALI-COs per heteroplasmic line and isogenic controls from different batches. Statistical analysis: one-way ANOVA with Tukey’s post hoc test (overall ANOVA p-values indicated in the graphs); **F.** Pseudotime analysis of scRNA-seq data, with branches representing the color-coded trajectory of cell-subtype specification (upper) or cluster identities (lower); **G.** Projection of transcriptomic data from heteroplasmic and control organoids onto a fetal brain reference dataset using scmap. The alluvial plot (left) shows cellular cluster mappings to human gestational ages. Bar plots represent the proportion of cells (middle) assigned to different fetal ages for each patient and isogenic control line. The rightmost bar plot illustrates neuronal cell type distributions mapped to fetal developmental stages; **H**. UMAP representation of cell cycle gene expression across all 11 identified cellular populations (left). The bar plot (middle) quantifies the proportion of cells in each cell cycle phase across heteroplasmic groups (low, medium, and high heteroplasmy). Statistical analysis: The Pearson’s chi-squared test was used to compare distributions of categorical data. P-values have been adjusted using the Benjamini-Hochberg procedure, suggesting no significant differences in cell cycle distribution across heteroplasmic organoids.

We next evaluated the tissue architecture by immunostaining for upper-layer (SATB2) and deep-layer (CTIP2) neuronal markers (Figure 2D and E and Figure S2D). Although the overall laminar organization was preserved in all organoids, evident from the segregation of deep-layer from upper-layer neurons, the proportion of CTIP2-positive cells was significantly reduced in high-heteroplasmy organoids. In contrast, SATB2-positive populations remained unaffected (Figure 2E). These findings indicate that high m.3243A>G heteroplasmy selectively compromises deep-layer neuronal populations.

Finally, we directly assessed whether the m.3243A>G heteroplasmy load affects cell type specification, maturation and cell cycle, potentially influencing the readout on m.3243A>G-driven mechanisms in neuronal pathology. To do so, we analysed the pseudotime trajectories in the control and heteroplasmic organoids using Monocle3^20^. This illustrated very similar transitions from radial glia (RG) towards mature DLN and ULN cell types following the typical developmental sequences in all groups (Figure 2F and Figure S2E). We then compared the transcriptomic maturity profiles of isogenic controls and heteroplasmic organoids with an age-specific reference dataset deriving from human fetal brain tissue^21^ using scmap^22^ (Figure 2G and Figure S2F). The maturity profiles across all samples aligned well with fetal brain cell transcriptomic signatures at 21 gestational weeks (140 days), including neurons, which closely matched the in vitro age of organoids (Figure 2G and Figure S2F). In addition, the cell cycle signatures inferred from merged transcriptomic data, revealed that most cells in the various heteroplasmic organoids were uniformly in the G1 phase, characteristic of post-mitotic neurons, while some cells were in the S and G2 phases, corresponding with progenitor cell identities (Figure 2H). Our findings therefore indicate that the m.3243A>G variant does not cause cell differentiation changes or cell cycle arrest, allowing accurate elucidation of mutation-driven pathologies in mature cortical neuronal populations.

### Cortical organoids with the m.3243A>G variant display a mitochondrial defect

To determine how varying heteroplasmy levels influence mitochondrial function in organoids we first examined OXPHOS protein abundances (Figure 3A). Organoids with high heteroplasmy exhibited reduced levels of NDUFB8, a Complex I subunit, whereas MTCO1, a Complex IV subunit, was decreased also in medium and high heteroplasmy organoids (Figure 3A,B). In contrast, SDHB (Complexes II) and ATP5A (Complex V) remained unchanged in all organoids (Figure S3A). These results confirm that the extent of OXPHOS disruption depends on heteroplasmy levels, with Complex I and IV being more susceptible to elevated mutation burden.

**Figure 3.**
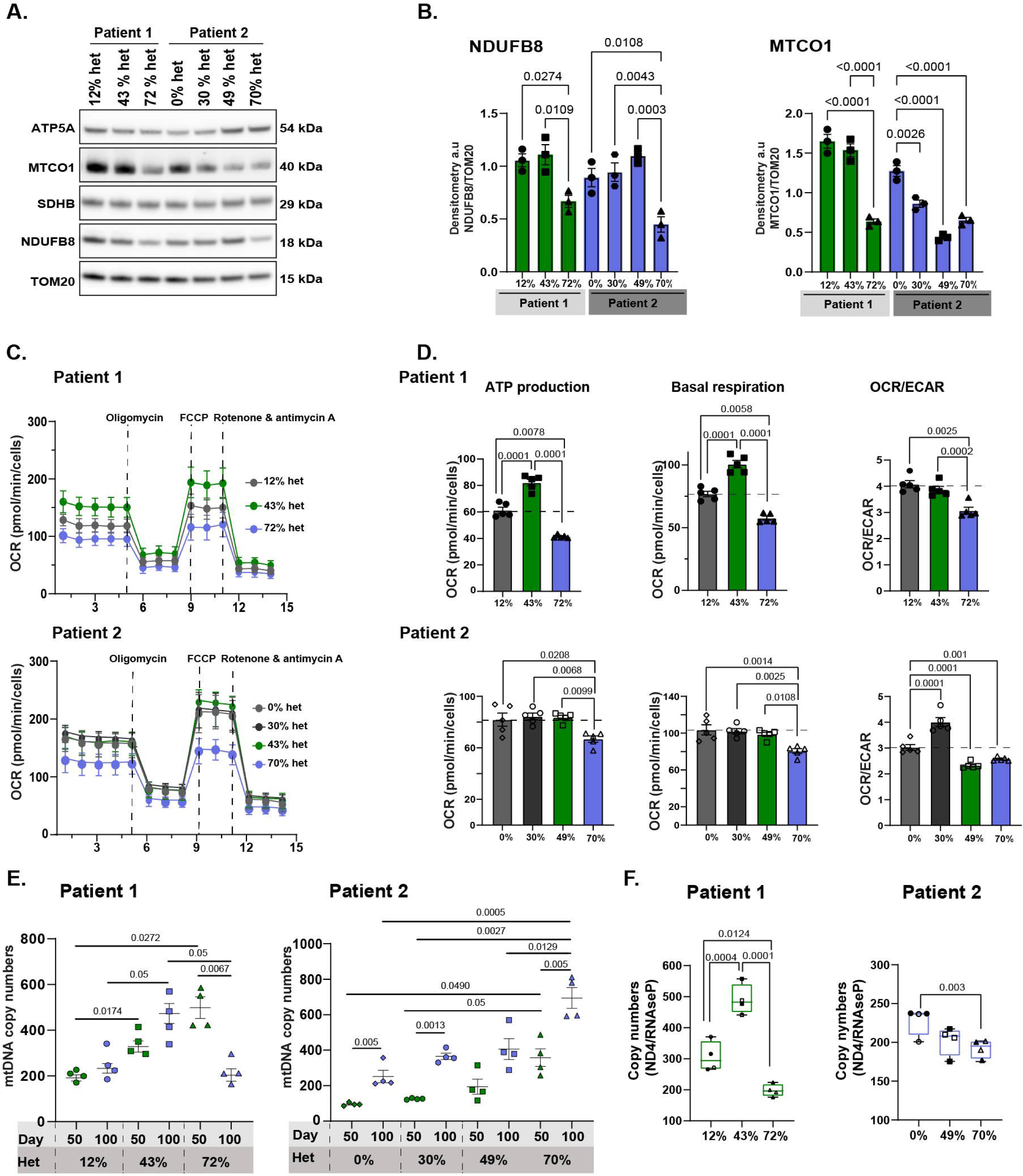
High heteroplasmy m.3243A>G organoids display a mitochondrial defect. **A.** Representative western blot (WB) images for ATP5A, MTCO1, SDHB, NDUFB8, TOM20, and ACTB. Corresponding molecular weights are displayed on the right side of the blot; **B.** Quantification of WB data for NDUFB8 (Complex I) and MTCO1 (Complex IV). Data were normalized to TOM20 to account for mitochondrial content. Data represent the mean ± SEM; *n* = 3 ALI-COs from three independent batches, with two organoids pooled per sample. Statistical analysis: one-way ANOVA with Tukey’s post hoc test (overall ANOVA *p*-values indicated in the graphs); **C.** Oxygen consumption rate (OCR) measurements in 100 DIV dissociated organoids. The upper panel represents data from Patient 1, and the lower panel represents data from Patient 2. OCR was measured at basal levels, followed by supplementation with 1 µM oligomycin, 1 µM FCCP, and 1 µM rotenone/antimycin A. Raw OCR values were normalized to total cell numbers, determined by Hoechst staining; **D.** Quantification of mitochondrial respiration. Bar plots show ATP-linked respiration (left), basal respiration or OCR (middle), and the OCR/ECAR ratio (right) to determine metabolic reliance on oxidative phosphorylation (OXPHOS) or glycolysis. The upper panel represents data from Patient 1, and the lower panel represents data from Patient 2. *n* = 5 dissociated ALI-COs for heteroplasmic and control organoids, with each experiment performed in triplicate to confirm trends. Statistical analysis: one-way ANOVA with Tukey’s post hoc test (overall ANOVA *p*-values indicated in the graphs); **E.** Dot plot showing individual measurements of mtDNA copy number in whole organoids at 50 and 100 DIV. Data represent the mean ± SEM of *n* = 4 ALI-COs from different batches. Statistical analysis: two-way ANOVA with Tukey’s post hoc test for multiple comparisons (overall ANOVA *p*-values indicated in the graphs); **F.** Box plots showing absolute quantification of mtDNA in neurons sorted from 100 DIV organoids. *n* = 2 independent sorting experiments from three organoids pooled per sample. Statistical analysis: one-way ANOVA with Tukey’s post hoc test (overall ANOVA *p*-values indicated in the graphs).

Real time metabolism was followed by the Seahorse assay, which was performed on cells dissociated from organoid slices. High heteroplasmy organoids showed a diminished capacity to generate energy through oxidative phosphorylation and are less adaptable to stress, as we observed reduced basal respiration, ATP production, maximal respiration, and spare respiratory capacity compared to medium and low heteroplasmy samples (Figure 3C and D, Figure S3B, C). High heteroplasmy levels were also associated with a tighter coupling of oxidative phosphorylation and a shift toward glycolysis, evident by a decreased proton leak and a lower OCR/ECAR ratio (Figure 3C and D, Figure S3B, C). In contrast, low and medium heteroplasmy organoids did not exhibit these respiration defects, as the mutation load remains below the pathogenic threshold that can cause mitochondrial dysfunction (Figure 3C and D, Figure S3C).

Next, we examined how cortical organoids regulate mtDNA content in response to metabolic dysfunction at 50 and 100 DIV. At 50 DIV, medium and high heteroplasmy organoids have increased mtDNA copy numbers compared to controls, consistent with a compensatory response to maintain mitochondrial function (Figure 3E). As organoids mature and accumulate more post-mitotic cells, low heteroplasmy organoids continued to increase mtDNA copy numbers between 50 and 100 DIV, with all medium and P2 high heteroplasmy organoids following a similar pattern (Figure 3E). In contrast, P1 high heteroplasmy (72%) organoids failed to sustain mtDNA expansion at 100 DIV, suggesting a threshold beyond which further amplification is not possible.

We examined mtDNA copy number regulation specifically in post-mitotic cells by isolating neurons and astrocytes from dissociated 100 DIV organoids using fluorescence-activated cell sorting (FACS) (Figure 3SD). In high heteroplasmy neurons mtDNA copy numbers were significantly reduced compared to controls (40% in P1 and 26% in P2), whereas medium heteroplasmy neurons exhibited more variability, suggesting the presence of compensatory mechanisms (Figure 3F). In astrocytes, mtDNA copy numbers decreased both in medium and high heteroplasmy conditions, with a significant reduction (43%) observed only in high heteroplasmy P1 astrocytes (Figure 3SE). Importantly, heteroplasmy levels in neurons and astrocytes remain unchanged and are comparable to those observed in the whole organoid (Figure S3F). Together all these findings highlight a critical heteroplasmy threshold beyond which mitochondrial compensation fails, leading to metabolic dysfunction and cell type-specific vulnerability.

### m.3243A>G drives transcriptomic signatures of perturbed neuronal homeostasis

By analysing differentially expressed genes (DEGs) from scRNA-seq data, we examined how mitochondrial dysfunction drives cell type specific changes in organoids. We compared transcriptomic signatures between high (72% het P1, 70% het P2) and low heteroplasmy (12% het P1, 0% het P2, 30% het P2) and between medium (43% het P1, 49% het P2) and low heteroplasmy. In both comparisons, intermediate progenitors consistently ranked among the top three populations with the highest number of DEGs. A striking difference emerged amongst neuronal populations. Deep-layer neurons (DLN5A and DLN5B) were most affected in high heteroplasmy, while DLN5A cells and developing upper-layer neurons (ULN) were the predominantly involved population in medium heteroplasmy (Figure 4A, B).

**Figure 4.**
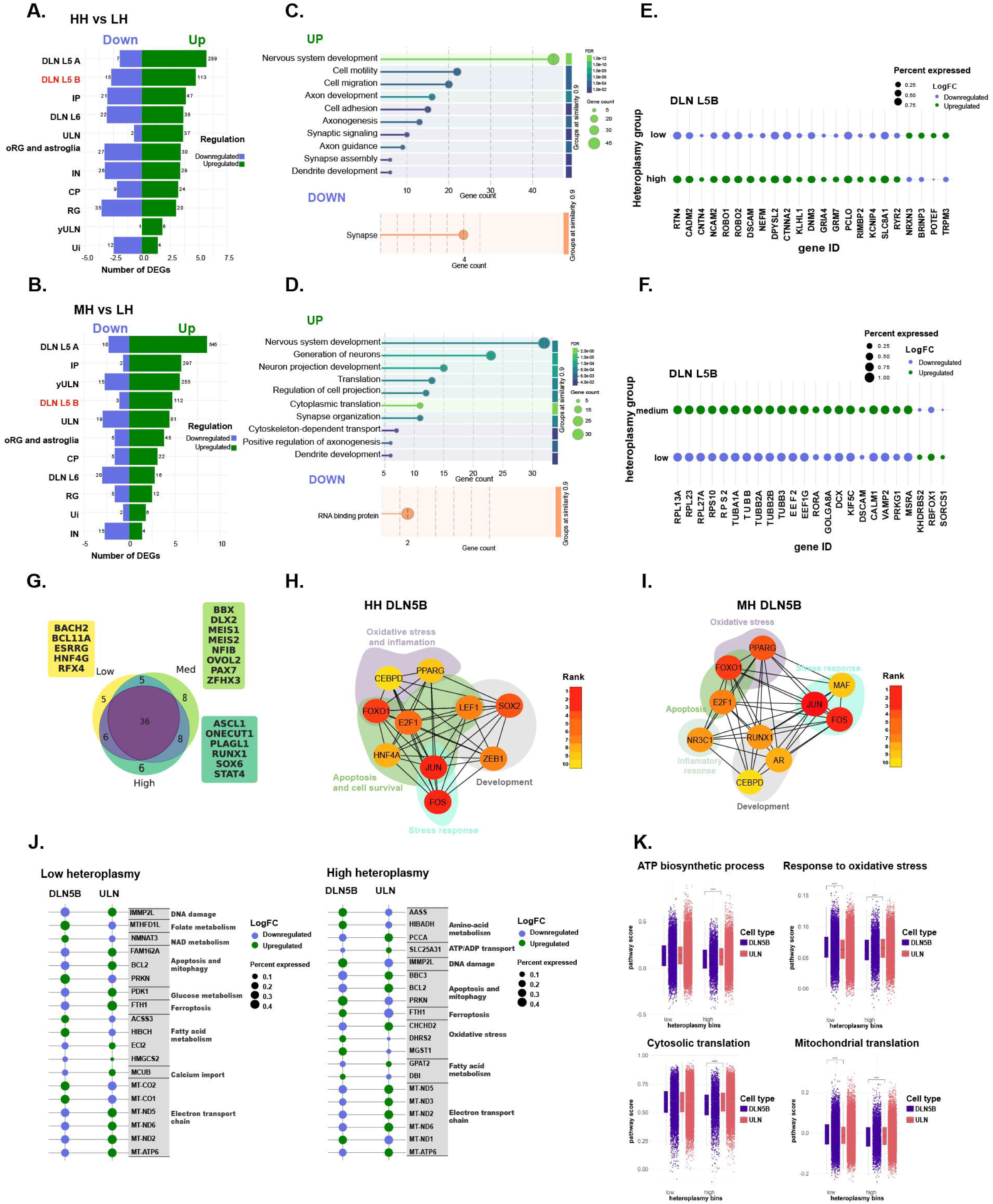
The m.3243A>G mutation impacts neuronal homeostasis A, **B**. Number of differentially expressed genes (DEGs) per cell type in high heteroplasmy (HH) vs. low heteroplasmy (LH) and medium heteroplasmy (MH) vs. low heteroplasmy (LH) (three slices per organoid line); **C, D.** Dot plots representing gene set enrichment analysis (GSEA) identifying the top 10 Gene Ontology (GO) terms with the largest term size enriched in DLN5B for both comparisons, performed using STRING^43^. Terms were grouped by similarity, with bubble color and connecting lines indicating the false discovery rate (FDR) value, while bubble size represents the number of enriched terms per pathway; **E, F.** Dot plots showing selected DEGs associated with the enriched GO terms identified in (C, D) for DLN5B in both comparisons; **G.** Venn diagram illustrating shared and unique transcription factors (TFs) identified in high, medium, and low heteroplasmy organoid groups. TF analysis was performed using SCENIC on merged data, representing transcription factors across all cell populations; **H, I**. Core network interaction maps (STRING) of transcription factors with predicted activity, identified using SCENIC. TFs were ranked by Maximal Clique Centrality (MCC) network topology (CytoHubba from Cytoscape), and functional domains were assigned based on gene/protein function (UniProt, https://www.uniprot.org). Node (circle) size corresponds to the MCC score, while color gradient represents MCC score ranking; **J.** Dot plots showing all DEGs mapping to MitoCarta identified in our single-cell RNA sequencing (scRNA-seq) analysis when comparing DLN5B with ULN in both low and high heteroplasmy conditions. Transcripts were categorized into functional domains based on gene/protein function (UniProt, https://www.uniprot.org); **K.** Box plots with individual data points representing GO enrichment of major pathways in DLN5B vs. ULN for both low and high heteroplasmy. The Wilcoxon test was used to compare the distribution of continuous variables between groups, such as gene expression levels.

Gene Ontology (GO) analysis revealed that high heteroplasmic DLN5B neurons show significant alterations in pathways related to cell adhesion, axon/dendritic development (*RTN4, CADM2, CNTN4, NCAM2, ROBO1/2, DSCAM*), and cytoskeletal remodelling (N*EFM, DPYSL2, CTNNA2, KLHL1, DNM3*), suggesting axonal disruption and degeneration. The upregulation in gene expression of glutamate receptor subunits (*GRIA4, GRM7*), presynaptic scaffolding proteins (*PCLO, RIMBP2*), and expression changes in ion channel (*KCNH7, KCNIP4*) and calcium exchanger genes (*SLC8A1, RYR2*) further indicate synaptic dysfunction and altered neuronal excitability (Figure 4C and E). In contrast, in medium heteroplasmy DLN5B neurons exhibited a stress-adaptive response, characterized by an enrichment of cytosolic translation related transcripts, including ribosomal proteins (*RPL13A, RPL23, RPL27A, RPS10, RPS2*) and elongation factors (*EEF2, EEF1G*). We observed an increased expression of tubulin genes (*TUBA1A, TUBB, TUBB2A, TUBB2B, TUBB3*) suggesting active cytoskeletal reorganization that supports axonal and dendritic development (Figure 4D and F).

Transcription factor (TF) inference analysis^23^ across the merged data identified 36 common TFs linked to organoid development and maturation (Figure S4A). In high heteroplasmy, we identified six uniquely expressed TFs, including *PLAGL1* and *STAT4*, which are associated with apoptosis and inflammation supporting a pathogenic environment. Medium heteroplasmic cells, express unique TFs predominantly involved in developmental pathways and cell maintenance, implying engagement of adaptive mechanisms to preserve neuronal function and regional identity (Figure 4G). Further TF activity analysis in high and medium heteroplasmic DLN5B cells using STRING networks and CytoHubba Maximal Clique Centrality analysis (embedded in Cytoscape)^24^ revealed a core set of stress-responsive TFs (*PPARG, FOXO1, E2F1, JUN, FOS*). High heteroplasmic neurons uniquely express additional TFs (*LEF1, HNF4A, ZEB1*) that reinforce apoptotic and oxidative stress signalling, whereas medium heteroplasmy neurons engage factors such as *NR3C1* and *PPARG* that can promote stress resilience (Figure 4H and I and Figure S4B).

To explore the potential causes for the cell stress changes predicted predominantly for DLN5B neurons, we compared their transcriptomic profiles with those of ULN neurons in control and high heteroplasmy cells. Control DLN5B neurons displayed a more oxidative metabolic profile with increased Complex IV transcripts (*MT-CO1, MT-CO2*), lower Complex I transcripts (*MT-ND2, MT-ND4, MT-ND6*) and enriched fatty acid metabolism, compared to ULN neurons, which are enriched for glycolysis-related GO terms (Figure 4J, K and Figure S4C, D). High heteroplasmy DLN5B neurons displayed metabolic disruptions compared to ULN, with a significant downregulation of ATP biosynthesis pathways together with decreased response to oxidative stress and increase in ferroptosis (*FTH1*), and DNA damage markers (*IMMP2L*) suggesting increased cellular stress and increased apoptosis. Additionally, mitochondrial and cytosolic translation-related GO terms were significantly overrepresented in high heteroplasmy DLN5B neurons compared with ULN, indicating cell type-specific translational dysfunction (Figure 4K). Together, these findings demonstrate that DLN5B neurons are more vulnerable to mitochondrial dysfunction, oxidative stress, and apoptotic signalling under high heteroplasmy compared to both low heteroplasmy conditions and high heteroplasmy ULN neurons, aligning with their predominant loss in patient-derived organoids.

### m.3243A>G high heteroplasmy promotes apoptosis of deep layer neurons

To extend our organoid findings to human pathology, we examined postmortem brain tissue from a 70-year-old male MELAS patient carrying the m.3243A>G variant (27% heteroplasmy detected in blood). The patient initially presented with mitochondrial myopathy, ophthalmoparesis, and fatigable weakness, with muscle biopsy showing abnormal myofibers, COX-/SDH-positive fibers, and selective type II fiber atrophy (Figure S4A). Following the development of acute neurological symptoms and SLEs, the patient died within 4 months. Brain autopsy revealed severe neuronal loss in the occipital lobe and marked glial fibrillary acidic protein (GFAP) immunoreactivity, suggesting glial cell responses (Figure S4B). Specifically, layer V, identified by its dense DAPI staining, showed a marked reduction in MAP2 immunoreactivity and the lack of cells with deep pyramidal neuronal morphology in the MELAS tissue (Figure 5A, white arrows) when compared to the control cortical samples. The upper layer also showed some decrease in MAP2^+^ neurons while layer VI appeared relatively preserved. Importantly, these findings well reflect the loss of CTIP2^+^ deep layer neurons found in cortical organoids derived from MELAS patients (Figure 2D, E), suggesting a shared mitochondrial mechanism of neuronal loss between the human brain tissue and cortical organoids.

**Figure 5.**
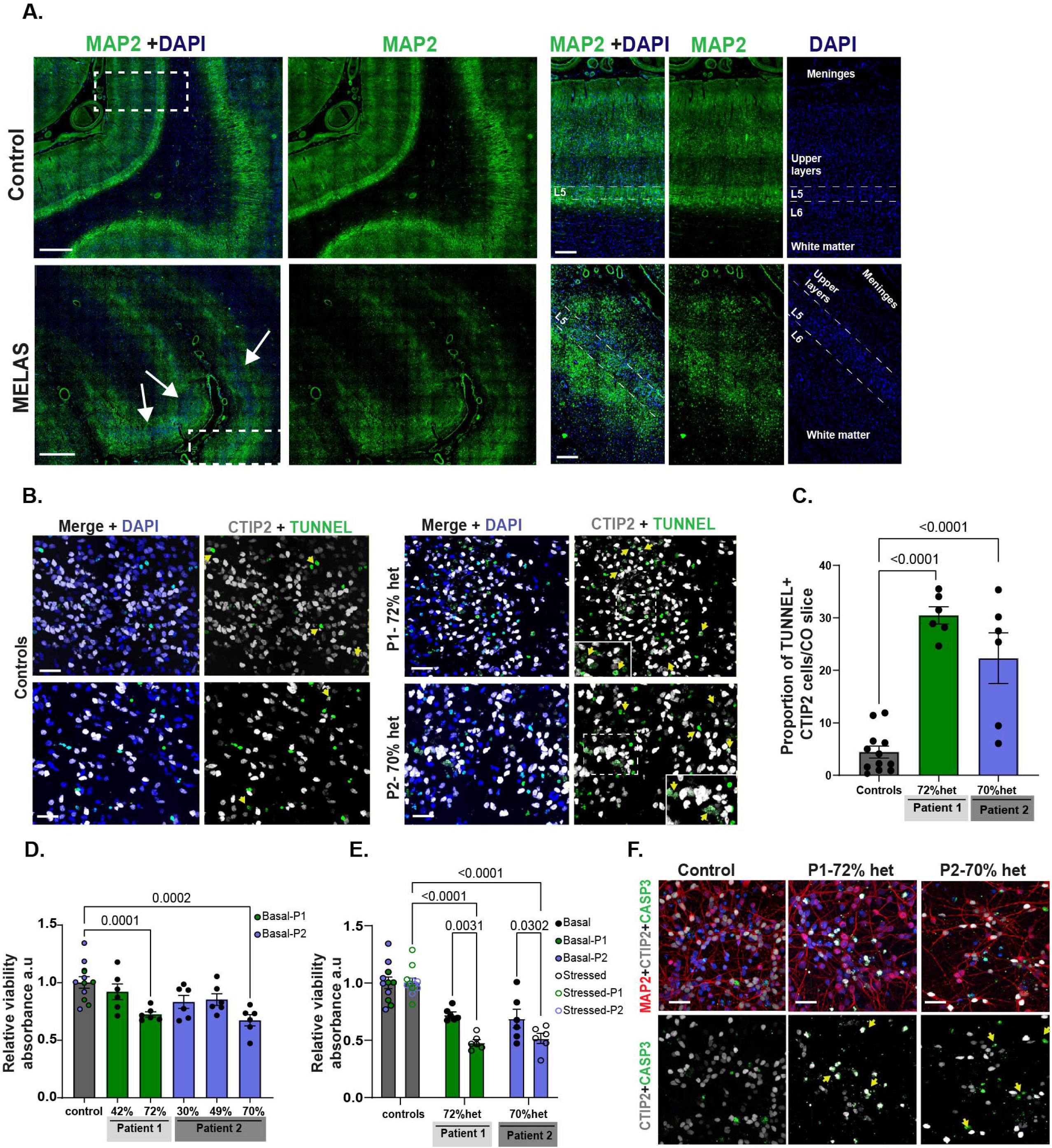
High heteroplasmy m.3243A>G mutation leads to neuronal apoptosis in deep layer neurons. **A.** Representative immunofluorescence images of human brain tissue from individuals with the m.3243A>G mutation and age-matched controls, stained for MAP2 (microtubule-associated protein 2, green), a neuronal marker. DAPI (blue) was used to label nuclei. Images illustrate neuronal morphology and MAP2 distribution in affected and control samples, with dotted white lines in the right panels marking layer 5 neurons. *n = 1*. Scale bar = 200 µm (right panel) and 50 µm (left panel); **B.** Representative immunofluorescence images of 100-day in vitro (DIV) cortical organoids with high heteroplasmy and isogenic controls, showing CTIP2LJ cells (grey) and TUNELLJ nuclei indicating apoptosis. Scale bar = 100 µm; **C.** Bar plots quantifying the proportion of TUNELLJ cells co-localizing with CTIP2, assessing apoptosis in deep-layer neurons. Data represent the mean ± SEM; *n = 6 independent ALI-COs* per heteroplasmic line and isogenic controls. Statistical analysis was performed using one-way ANOVA with Dunnett’s post hoc test, with overall ANOVA p-values indicated in the graphs; **D.** Bar chart showing relative cell viability measured by the PrestoBlue assay in 2D-derived deep-layer neurons 30 DIV under basal conditions. Data represent the mean ± SEM; *n = 6* from three independent batches of neurons. Statistical analysis was performed using one-way ANOVA with Dunnett’s post hoc test, with overall ANOVA p-values indicated in the graphs; **E.** Bar chart showing relative cell viability determined by the PrestoBlue assay under basal and stressed conditions, where mitochondrial function was acutely stimulated using 10 µM GSK-2837808A + 0.1% AlbuMAX for 12 hours. Data represent the mean ± SEM; *n = 6* from three independent batches of neurons. Statistical analysis was performed using two-way ANOVA with Tukey’s post hoc test for multiple comparisons, with overall ANOVA p-values indicated in the graphs; **F.** Representative immunofluorescence images of CTIP2+ cells (grey), cleaved CASP3 (green) and MAP2 (red) in isogenic control and high heteroplasmy glutaminergic 2D neurons. n = 3 biological replicates. Scale bar = 50 µm.

We next determined whether the neuronal loss is a result of apoptotic cell death in MELAS organoids. Apoptosis was assessed in organoids with high heteroplasmy and isogenic controls using a terminal deoxynucleotidyl transferase dUTP nick end labelling (TUNEL) assay (Figure 5B). We found an increased number of TUNEL-positive cells colocalizing with CTIP2 immunoreactivity, indicating that apoptosis contributes to the reduced DLN5 population in high heteroplasmic organoids (Figure 5B and C). To further explore the contribution of mtDNA heteroplasmy-related cell death in this specific neuronal population, we generated CTIP2^+^ DLN5 neuronal mono-cultures obtained from a homogeneous pool of neuronal progenitors (Figure S5C, D and E). By day 30, high heteroplasmy neurons showed decreased viability compared with medium and low heteroplasmy cells (Figure 5D). The increased immunoreactivity of cleaved CASPASE3 in CTIP2 neurons supported their apoptotic cell death (Figure 5F). LDHA blockade evoked metabolic stress and lipid supplementation driven induction of mitochondrial function revealed further decrease in viability in high heteroplasmy neurons, while medium and low heteroplasmy cells remained mostly unaffected (Figure 6E, Figure S5F). These findings corroborates that the death of high heteroplasmy DLN5B neurons is exacerbated by mitochondrial stress.

**Figure 6.**
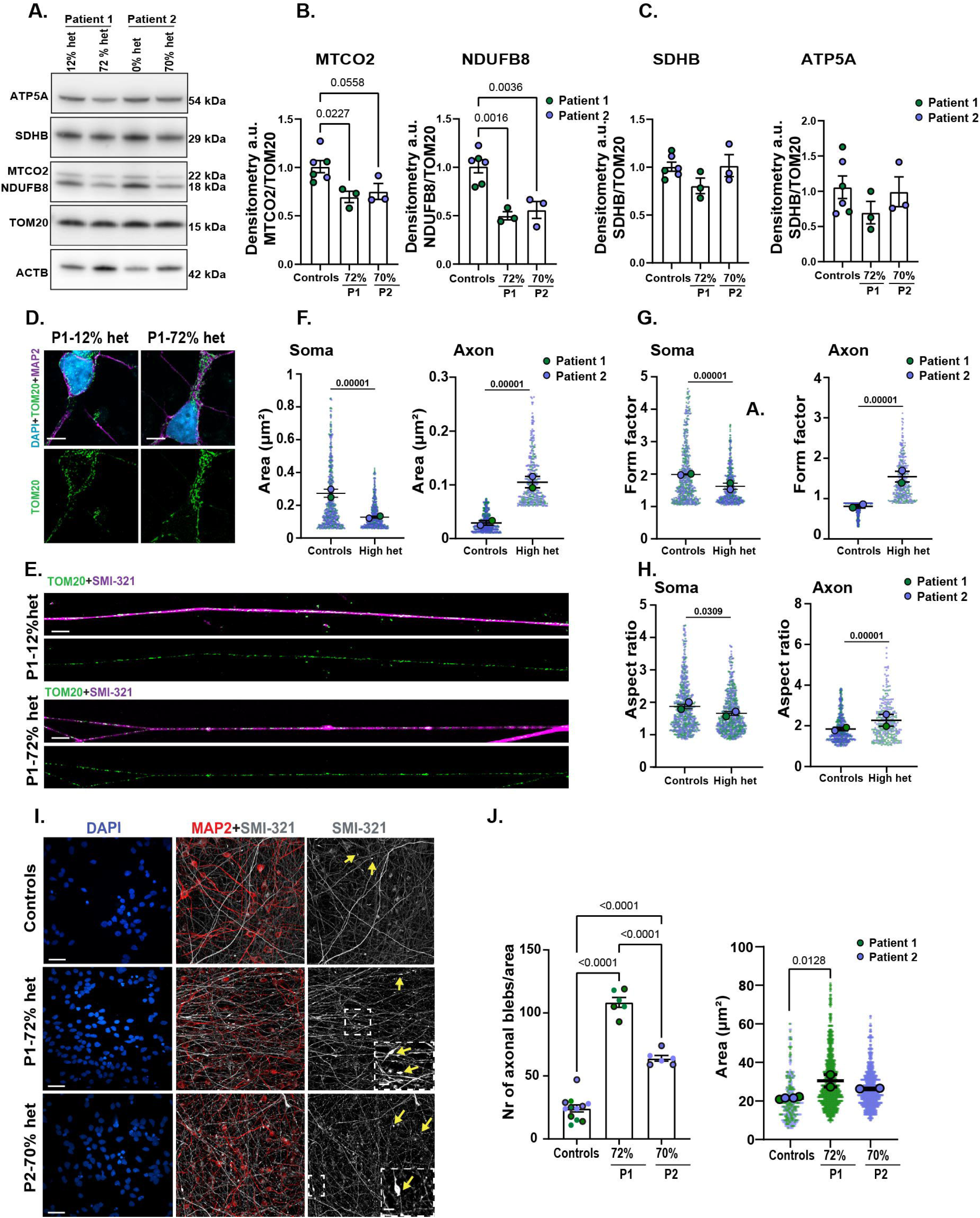
Deep layer neurons display mitochondrial defect. **A.** Representative WB images for ATP5A (Complex V), SDHB (Complex II), MTCO2 (Complex IV), NDUFB8 (Complex I), TOM20, and ACTB in 2D 30-day in vitro (DIV) glutamatergic neurons. Corresponding molecular weights are displayed on the right side of the blot; **B, C.** Quantification of Western blot data for MTCO2 (Complex IV), NDUFB8 (Complex I), SDHB (Complex II), and ATP5A (Complex V). Protein levels were normalized to TOM20 to account for mitochondrial content, while ACTB was used as a loading control. Data represent the mean ± SEM; *n = 3* from independent batches. Statistical analysis was performed using one-way ANOVA with Tukey’s post hoc test, with overall ANOVA p-values indicated in the graphs; **D.** Representative immunofluorescence images showing MAP2 (magenta) marking dendrites and cell bodies, and TOM20 (green) marking mitochondria in 2D control and high-heteroplasmy neurons. Scale bar = 50 µm; **E.** Representative immunofluorescence images of axonal mitochondria labeled with TOM20 (green) and the axonal neurofilament marker SMI-321 (magenta) in 2D cortical neurons from controls and high-heteroplasmy lines. Scale bar = 50 µm; **F-H.** Dot plots representing mitochondrial morphology quantification in soma and axons, assessing mitochondrial area, form factor, and aspect ratio. Larger dot plots indicate the median of all quantified regions of interest (ROIs) for each line. Data represent the mean ± SEM; *n = 4*, with two control lines and two high-heteroplasmy lines derived from two independent batches. Statistical analysis was performed using one-way ANOVA with Dunnett’s test, with overall ANOVA p-values indicated in the graphs; **I.** Immunofluorescence images showing axons (SMI-321, gray) segregating from dendrites (MAP2, red) in 2D cortical neurons from control and high-heteroplasmy lines. Insets and arrows highlight axonal fragmentation and blebs. Scale bar = 50 µm; **J.** Bar plots quantifying the number of axonal blebs per area, where dots represent the number of regions of interest (left panel) and the area of axonal blebs (right panel) for images shown in (I). Data represent the mean ± SEM; *n = 4*, with two control lines and two high-heteroplasmy lines derived from two independent batches. Statistical analysis was performed using one-way ANOVA with Dunnett’s test, with overall ANOVA p-values indicated in the graphs.

### High heteroplasmy alters mitochondrial morphology and axon integrity in deep layer neurons

To explore further mitochondrial abnormalities in deep layer neurons, we compared OXPHOS protein levels and mitochondrial morphology in cortical neuronal mono-cultures with high heteroplasmy versus isogenic control cells. Western blotting revealed a consistent reduction in MTCO1 (Complex IV) and NDUFB8 (Complex I) in high heteroplasmy deep-layer neurons, while SDHB (Complex II) and ATP5A (Complex V) remained largely unchanged compared to controls (Figure 6A-C). These results align with our findings revealed in cortical organoids (Figure 3A, B and Figure 6A, C), supporting the mitochondrial defects in high heteroplasmy neurons.

To determine whether these molecular alterations in OXPHOS were accompanied by structural changes, we examined mitochondrial morphology in DLN using TOM20 as a mitochondrial marker, MAP2 for dendrites (cell body), and SMI-32 for axons (Figure 6D–H and Figure S5G). In the soma, mitochondria in high heteroplasmic neurons displayed reduced area, indicative of smaller, fragmented mitochondria (Figure 6D and F). This was accompanied by changes in form factor and aspect ratio, further supporting abnormal mitochondrial morphology (Figure 6D-H). In contrast, axonal mitochondria in high heteroplasmy neurons displayed increased area, particularly accumulating in axonal blebs (white arrows), alongside elevated form factor and aspect ratio (Figure 6E-H) suggesting a more elongated mitochondrial morphology.

The abnormal mitochondrial morphology and axonal distribution, along with scRNA-seq data indicating axonal dysfunction, in high heteroplasmy deep layer neurons (Figure 4C), prompted us to examine axonal structure. Immunofluorescence of SMI-32, an axonal label, revealed significantly increased number of axonal blebs, with enlarged bleb area compared to controls (Figure 6I, J). Our findings indicate that mitochondrial dysfunction in high heteroplasmy neurons contributes to axonal structural abnormalities with signatures of early degeneration, which highlights a mitochondrial stress-induced axonal vulnerability in deep layer neurons.

## DISCUSSION

Here, we developed a 3D human cortical brain organoid model carrying the heteroplasmic m.3243A>G mtDNA mutation, the most common pathogenic mtDNA variant^25^. So far, one recent study used a human 3D brain tissue model harbouring a nuclear mutation in *SURF1*^26^ to study mitochondrial dysfunction, while cortical organoid platforms have not been utilized to explore mtDNA mutations^16^. The use of these advanced human systems is a pre-requisite for studying common mtDNA mutations due to the lack of animal models. Importantly, organoid slice culture preparations allow long term cultures, cell type diversity and maturation, improving their reliability to model human diseases^17,19,27^.

Our 3D human cortical brain organoid slice model of m.3243A>G successfully recapitulated mature neuronal phenotypes while exhibiting distinct molecular pathological characteristics of mitochondrial dysfunction. In this study, we uncovered key molecular mechanisms underlying MELAS, the most severe phenotype of m.3243A>G observed in patients, and identified other brain related neuronal pathologies associated with this common mtDNA mutation. Using isogenic and mutant cortical organoids derived from seven iPSC clones from two patients with low, medium, and high heteroplasmy levels, we systematically examined how different mutation burdens affect neuronal homogeneity, maturity, and diversity, as well as their impact on neuronal cell homeostasis.

Heteroplasmy levels in our organoids remained stable throughout their formation, showing consistency across different organoids and batches, suggesting no major selective pressure for or against the mutation. Likewise, neuron- and astrocyte-specific mutation loads were comparable, suggesting that heteroplasmy remains uniform across different mature cell populations within the organoids. This stability aligns with previous studies indicating that rapid shifts in heteroplasmy primarily occur during the reprogramming of fibroblasts into iPSCs, whereas heteroplasmy levels remain relatively stable throughout neuronal development^12,13^. In human post-mortem studies, brain heteroplasmy levels of the m.3243A>G mutation have been reported to be high in both symptomatic and asymptomatic individuals, suggesting that while heteroplasmy contributes to the clinical manifestations of MELAS, it is not the sole determinant of disease severity^28,29^. Additional factors such as mtDNA copy number, nuclear genetic background, and environmental influences likely play critical roles in modulating the phenotype^9^. These findings highlight the complexity of mitochondrial disease and suggest that stable heteroplasmy in organoid models provides a useful system for studying cell type-specific mitochondrial dysfunction without the confounding effects of dynamic heteroplasmy shifts.

Our findings demonstrate that high heteroplasmy levels do not affect cellular composition or heterogeneity, as all 10 identified neuronal populations with single-cell RNA sequencing, were present in both control and m.3243A>G organoids. These organoids also contain mature neuronal subtypes, including distinct deep- and upper-layer cortical architecture. Furthermore, no evidence of cell cycle arrest was observed in high heteroplasmy organoids compared to medium or low heteroplasmy. While mitochondrial dysfunction can influence neuronal maturation^30,31^, our results suggest that these effects may be more subtle in m.3243A>G disease or only become apparent at higher heteroplasmy thresholds (<80% het). This aligns with the milder and later onset manifestation of MELAS in adults^9,32^, in contrast to severe early childhood-onset mitochondrial diseases such as Leigh Syndrome due to *SURF1* mutations, where brain organoids showed that morphogenesis is profoundly disrupted^26^.

Mitochondrial dysfunction correlated with heteroplasmy load in cortical organoids, with high-heteroplasmic organoids showing the most severe OXPHOS defects. Complex I expression was consistently reduced, while complex IV levels varied across heteroplasmy loads, mirroring findings from m.3243A>G patient brain autopsies^29^. Biochemical assays confirmed metabolic impairment in high-heteroplasmy organoids with limited ability to adapt to metabolic stress. This vulnerability was further supported by studies on 2D neurons, where increased mitochondrial activity and stress led to decreased viability, likely due to apoptosis from excessive metabolic burden. In contrast, medium heteroplasmy organoids with normal mitochondrial function and one high heteroplasmy organoid (P2-70%), exhibited increased mtDNA copy numbers, suggesting a compensatory mechanism to sustain OXPHOS by preserving wild-type mtDNA. This effect appears limited beyond a certain threshold, leading to irreversible mitochondrial defects. Despite similar heteroplasmy levels in high heteroplasmy P1 and P2 at 100 DIV, their distinct mtDNA trajectories suggest that heteroplasmy levels have a strong influence on mtDNA expansion at earlier stages. A similar adaptation has been observed in dopaminergic neurons, where increased mtDNA copy number have been proposed as a compensatory response to age-related mitochondrial dysfunction^33^. Importantly, since heteroplasmy levels are similar between neurons, astrocytes, and whole organoids copy number alterations are unlikely to be the main factor driving the mtDNA differences between different cell types and organoid preparations. This suggests that mtDNA maintenance is regulated in a cell type-specific manner, with distinct mechanisms operating in post-mitotic and proliferative cells.

Another important finding of our study was the predominant loss of deep-layer neurons (DLNs) in high-heteroplasmy m.3243A>G organoids, while upper-layer neurons (ULNs) remained largely preserved. Our scRNA-seq analysis suggests that this vulnerability is linked to intrinsic transcriptional differences between these main neuronal cell types that make DLNs more prone to apoptosis, oxidative stress, and metabolic dysfunction. Specifically, DLNs exhibited dysregulation in pathways associated with cell adhesion, cytoskeletal remodelling, axon and dendritic homeostasis when compared to isogenic control neurons. These transcriptional changes were reflected in our functional assays, which showed increased apoptosis and axonal degeneration in DLNs, alongside alterations in axonal mitochondrial morphology. The increased susceptibility of DLNs to mitochondrial dysfunction can be attributed to their distinct anatomical and physiological properties. Amongst cortical projection neurons, *CTIP2*-expressing DLN5 neurons, extend long-range connections to the spinal cord, brainstem, and thalamic regions^34,35^. Maintaining such extensive networks requires high metabolic energy, partly due to their longer axons and depolarized resting membrane potentials. This increased bioenergetic demand likely makes DLNs particularly vulnerable to disruptions in ATP production and oxidative phosphorylation caused by the m.3243A>G mutation. Importantly, DLNs are also affected in other neurological disorders such as amyotrophic lateral sclerosis, Parkinson’s disease, Alzheimer’s, epilepsy, further supporting their inherent susceptibility to disease and pathological changes^17,36–39^.

In comparison, ULNs have shorter axons and primarily project local^40^, they were preserved in high heteroplasmic organoids, suggesting a greater resilience to mitochondrial dysfunction. Direct comparison between high heteroplasmic DLN5B and high heteroplasmic ULNs revealed that DLN5 neurons exhibited greater metabolic disruption, including downregulation of ATP biosynthesis pathways and mitochondrial/cytosolic translation-related transcripts. Additionally, DLN5B neurons showed elevated expression of oxidative stress markers, ferroptosis-associated genes, and DNA damage markers, indicating higher levels of mitochondrial stress and reduced cellular resilience compared to ULNs. These findings suggest that ULNs are more capable of withstanding mitochondrial dysfunction, potentially due to a more effective metabolic compensation mechanism or lower energy requirements. Together, our results highlight a crucial link between neuronal subtype-specific metabolic demands and their differential vulnerability to mitochondrial dysfunction.

To correlate our findings with brain pathology, we examined autopsy data from a patient who died of m.3243A>G-related SLEs and detected severe neuronal loss. Postmortem analyses of m.3243A>G patients have consistently revealed cortical lesions and laminar necrosis (often affecting layer III and V) in parietal, lateral temporal, and occipital cortices, leading to clinical manifestations such as cortical blindness and homonymous hemianopia^29^. These findings suggest a selective vulnerability of specific neuronal populations, mirroring the deep-layer neuronal loss observed in our organoid model. Patients with high heteroplasmic m.3243A>G often experience SLEs with fluctuating metabolic lesions in cortical-subcortical areas, commonly associated with complex epileptic activity and progressive brain atrophy^10^. Although advanced neuroimaging techniques such as 7 Tesla MRI and PET with mitochondrial ligands have been used to study MELAS patient brains^41,42^, these approaches lack the resolution to identify cell type-specific neuronal dysfunction. This underscores the need for organoid models, which provide a unique platform for dissecting the dynamic changes of mitochondrial pathology at the cellular level and can facilitate therapeutic development targeting specific neuronal populations.

A limitation of our study is that we primarily focused on neurons and not on non-neuronal cell types, such as glial cells that may also contribute to the phenotype observed in m.3243A>G patients. To fully understand their role, future studies should investigate glial dysfunction using older organoids (>150 DIV), which allow for more advanced glial maturation. Additionally, since m.3243A>G patients frequently present with seizures, incorporating more complex organoid models that include GABAergic interneurons will be essential for studying electrophysiological dysfunction. Such models, combined with multi-electrode array (MEA) recordings, will provide deeper insights into the mitochondrial aspects of epilepsy and neurodegeneration. Future studies integrating single-cell resolution approaches should help refine our understanding of neuronal and glial susceptibility to mitochondrial dysfunction.

Overall, our study provides a novel mechanistic insight into cell type-specific mitochondrial defects caused by m.3243A>G in the brain, revealing how neuronal vulnerability is shaped by metabolic stress and bioenergetic demands. By leveraging human brain organoids, we established a powerful platform for testing potential treatments, including emerging mtDNA editing technologies^6^. Beyond therapeutic development, understanding mitochondrial dysfunction at the cellular level is critical not only for mitochondrial diseases but also for neurodegenerative, neurodevelopmental and psychiatric disorders, where mitochondrial deficits contribute to disease progression. We anticipate that using these human platforms will help to understand how mitochondrial function affect neuronal resilience, and degeneration, which could open new avenues for targeted interventions.

## RESOURCE AVAILABILITY

### Materials availability

This study has not generated new, unique reagents. Further information and requests for resources and reagents should be directed and will be fulfilled by lead contacts Rita Horvath, Andras Lakatos and Denisa Hathazi.

### Data and code availability

Single cell RNA seq data has been deposited at the European-Genome-phenome Archive (EGA), which is hosted by EBI and the CRG (in progress)

High resolution scans of all blots presented in this paper have been included in Source Data to this manuscript.

This paper does not report any original code.

Data is available in the manuscript or the supplementary materials. Any additional information required to reanalyze the data reported in this work paper is available from the lead contact upon request.

## ACKNOWLEDGMENTS

D.H. is supported by the Guarantors of Brain Non-Clinical Postdoctoral Fellowship. A.L. is supported by the Medical Research Council (UK) Senior Clinical Fellowship Award (MR/X006867/1). R.H. is supported by the Wellcome Discovery Award (226653/Z/22/Z), the Medical Research Council (UK) (MR/V009346/1), the Hereditary Neuropathy Foundation, the AFM-Telethon, the Ataxia UK, the Action for AT, the Muscular Dystrophy UK, the Rosetrees Trust (PGL23/100048), the LifeArc Centre to Treat Mitochondrial Diseases (LAC-TreatMito) and the UKRI/Horizon Europe Guarantee MSCA Doctoral Network Programme (Project 101120256: MMM). She was also supported by an MRC strategic award to establish an International Centre for Genomic Medicine in Neuromuscular Diseases (ICGNMD) MR/S005021/1. This research was supported by the NIHR Cambridge Biomedical Research Centre (BRC-1215-20014). The views expressed are those of the authors and not necessarily those of the NIHR or the Department of Health and Social Care.

We would like to thank Huw Naylor, Dr. Andreas Bruckbauer, Dr. Fadwa Joud, Heather Zecchini at the Light Microscopy Core Facility from the Cancer Research UK Cambridge Institute for their assistance with microscopy and data analysis. We would like to thank Gabriela Grondys-Kotarba and Dr. Reiner Schulte from the Flow Cytometry Core Facility at the Cambridge Institute for Medical Research for their help with cell sorting. We would like to thank also the Genomics facility from the Cambridge Stem Cell institute for preparing the scRNA sequencing library preparation.

We would like to thank Dr. George Gibbons and Dr. Kornélia Szebényi for valuable discussion and advice on organoid culture. D.H. would also like to thank George Nolan for his kind guidance with staining human brain. We thank Valeria Chichagova for generating some of the iPSC clones derived from Patient 1.

## AUTHORS CONTRIBUTIONS

Conceptualization: D.H., A.L., R.H.; methodology and investigation: D.H. A.L., R.H, D.L, M.Z; human organoid cultures: D.H., J.M.; scRNA-seq analysis: C.L., P.C.; Seahorse: D.H., O.P.; Imaging: D.H., D.L.; imaging analysis: D.H., D.L., M.Z.M; post-mortem brain samples: A.K. and D.H.; cells: M.L., T.K., E.M.T, L.L; manuscript writing: D.H., A.L., R.H., manuscript reviewing: D.H., D.L., M.Z.M, J.M., P.C., A.L., R.H; funding acquisition: R.H.

## STAR METHODS

### KEY RESOURCES TABLE

| Reagent or resource | Source | Identifier |
| --- | --- | --- |
| **Antibodies** |  |  |
| ACTB | Proteintech | Cat# 20536-1-AP,<br>RRID:AB_10700003 |
| Cleaved CASP3 | Cell Signalling | Cat# 9664, RRID:AB_2070042 |
| CTIP2 | Abcam | Cat# ab18465,<br>RRID:AB_2064130 |
| FOXG1 | Abcam | Cat# ab18259,<br>RRID:AB_732415 |
| GAD2 | Cell Signalling | Cat# 5843, RRID:AB_10835855 |
| GFAP | Antibodies.com | Cat# A85307,<br>RRID:AB_2748894 |
| HEPACAM | R and D Systems | Cat# MAB4108,<br>RRID:AB_2117687 |
| L1CAM | R and D Systems | Cat# MAB7771,<br>RRID:AB_3658914 |
| MAP2 | Antibodies.com | Cat# A85363,<br>RRID:AB_2748940 |
| MTCO1 | Abcam | Cat# ab14705,<br>RRID:AB_2084810 |
| NEST | Sigma-Aldrich | Cat# N5413,<br>RRID:AB_1841032 |
| SATB2 | Abcam | Cat# ab51502,<br>RRID:AB_882455 |
| SMI-321 | BioLegend | Cat# 837904,<br>RRID:AB_2566782 |
| SOX2 | R and D Systems | Cat# AF2018,<br>RRID:AB_355110 |
| TOM20 | Abcam | Cat# ab186735,<br>RRID:AB_2889972 |
| Total OXPHOS Human WB<br>Antibody Coctail Kit | Abcam | Cat# ab110411,<br>RRID:AB_2756818 |
| TUJ | Abcam | Cat# ab78078,<br>RRID:AB_2256751 |
| Donkey Anti Goat<br>IgG(H+L) Highly Crossed<br>Absorbed Secondary<br>Antibody, Alexa Fluor Plus<br>647 | Thermo Fisher Scientific | Cat# A32849TR,<br>RRID:AB_2866498 |
| Donkey Anti Mouse<br>IgG(H+L) Highly Crossed<br>Absorbed Secondary<br>Antibody, Alexa Fluor 568 | Thermo Fisher Scientific | Cat# A10037,<br>RRID:AB_11180865 |
| Donkey Anti Rabbit<br>IgG(H+L) Highly-Cross<br>Absorbed Secondary<br>Antibody, Alexa Fluor 488 | Thermo Fisher Scientific | Cat# A-21206,<br>RRID:AB_2535792 |
| Donkey Anti Rabbit IgG(H+L) Highly-Cross Absorbed Secondary Antibody, Alexa Fluor 568 | Thermo Fisher Scientific | Cat# SA5-10042, RRID:AB_2556622 |
| Donkey Anti Rat IgG H&L Alexa Fluor 647 preabsorbed | Abcam | Cat# ab150155, RRID:AB_2813835 |
| Goat Anti Chicken IgG(H+L) Cross-Absorbed Secondary Antibody Alexa Flour 488 | Thermo Fisher Scientific | Cat# A-11006, RRID:AB_2534074 |
| Goat Anti Chicken IgY(H+L) Cross-Absorbed Secondary Antibody Alexa Flour 647 | Thermo Fisher Scientific | Cat# A48260, RRID:AB_2890271 |
| Goat Anti Mouse IgG (H+L) Secondary Antibody, HRP | Thermo Fisher Scientific | Cat# G-21040, RRID:AB_2536527 |
| Goat Anti Rabbit IgG (H+L) Secondary Antibody, HRP | Thermo Fisher Scientific | Cat# 31462, RRID:AB_228338 |
| **Chemicals, peptides and Recombinant proteins** |  |  |
| DAPI Nuclear Counterstain | Thermo Fisher Scientific | Cat# 62248 |
| Accutase | Sigma-Aldrich | Cat# A1110501 |
| Antibiotic-Antimycotic (100X) | Thermo Fisher Scientific | Cat# 15240062 |
| Antimycin a from Streptomyces sp. | Sigma-Aldrich | Cat# A8674-25MG |
| Apai, New England Biolabs, ApaI, Size=5,000 units | New England Biolabs (NEB) | Cat# R0114S |
| B-27™ Supplement (50X), minus vitamin A | Thermo Fisher Scientific | Cat# 12-587-010 |
| B-27™ Supplement (50X), serum free | Thermo Fisher Scientific | Cat# 17504044 |
| Bovine Serum Albumin, lyophilized powder, BioReagent, suitable for cell culture, | Sigma-Aldrich | Cat# A9418-100G |
| BrainPhys™ Neuronal Medium, Liquid | STEMCELL Technologies | Cat# 5790 |
| Complete™, Mini, EDTA-free Protease Inhibitor Cocktail | Sigma-Aldrich | Cat# 11836170001 |
| Compound E | STEMCELL Technologies | Cat# 73952 |
| Costar 24 Well Clear Flat Bottom, Ultra Low Attachment Multiple Well Plates with Lid, Sterile, Individually Wrapped, 24/Case | Sigma-Aldrich | Cat# CLS3473-24EA |
| Costar 96 Well Clear Round Bottom Ultra Low Attachment Microplate with Lid, Sterile, Individually Wrapped, 24/case | Sigma-Aldrich | Cat# CLS7007-24EA |
| Cytiva's Amersham ECL Select Western Blotting Detection Reagent | Cytiva | Cat# RPN2235 |
| DMSO | Sigma-Aldrich | Cat# D2650 |
| Dnase I,from bovine pancreas | Sigma-Aldrich | Cat# 11284932001 |
| Donkey serum | Sigma-Aldrich | Cat# D9663-10ML |
| Dpbs, calcium, magnesium | Thermo Fisher Scientific | Cat# 14-040-133 |
| Dpbs, no calcium, no magnesium | Thermo Fisher Scientific | Cat# 14-190-144 |
| Geltrex | Thermo Fisher Scientific | Cat# A1413302 |
| Gibco DMEM/F 12, GlutaMAX Supplement | Thermo Fisher Scientific | Cat# 10-565-018 |
| Glucose Solution | Fisher Scientific | Cat# A2494001 |
| GlutaMAX™ Supplement | Thermo Fisher Scientific | Cat# 35050061 |
| LDN193189 hydrochloride, ≥98% (HPLC) | Sigma-Aldrich | Cat# SML0559-5MG |
| L-Glutamine (200 mM) | Thermo Fisher Scientific | Cat# 25030024 |
| Lipofuscin Autofluorescence Quencher | Biotum | Cat# 23007 |
| Live/dead™ Fixable Near-IR Dead Cell Stain Kit, for 633 or 635 nm excitation | Thermo Fisher Scientific | Cat# L10119 |
| Matrigel Matrix hESC-Qualified Mouse 5 MI | Corning | Cat# 354277 |
| Molecular Probes ProLong Gold Antifade Mountant | Thermo Fisher Scientific | Cat# P36934 |
| mTeSR Plus | STEMCELL Technologies | Cat# 100-0276 |
| MyTaq Red DNA Polymerase, 500 Units | Meridian Life Science | Cat# BIO-21108 |
| N-2 Supplement (100X) | Thermo Fisher Scientific | Cat# 17502048 |
| Neurobasal™ Medium | Thermo Fisher Scientific | Cat# 21103049 |
| Neurobasal™ Medium, minus phenol red | Thermo Fisher Scientific | Cat# 12348017 |
| NeuroCult SM1 Without Vitamin A | STEMCELL Technologies | Cat# 5731 |
| NeuroCult™ SM1 Neuronal Supplement | STEMCELL Technologies | Cat# 5711 |
| Normal Goat Serum (ab7481) | Abcam | Cat# ab7481 |
| Oligomycin from Streptomyces diastatochromogenes, ≥90% total oligomycins basis (HPLC) | Sigma-Aldrich | Cat# O4876-5MG |
| PDS Kit, Papain Vial | Worthington Biochemical | Cat# LK003176 |
| Penicillin-Streptomycin (5,000 U/mL) | Thermo Fisher Scientific | Cat# 15070063 |
| PICM0RG50 \| Millicell Cell Culture Insert | Sigma-Aldrich | Cat# PICM0RG50 |
| Pierce™ Methanol-free Formaldehyde Ampules | Thermo Fisher Scientific | Cat# 28908 |
| Pierce™ Rapid Gold BCA Protein Assay Kit | Thermo Fisher Scientific | Cat# A53225 |
| ProLong® Gold antifade reagent with DAPI | Invitrogen | Cat# P36931 |
| Recombinant Human BDNF | PeproTech | Cat# 450-02 |
| Recombinant Human FGF-basic (154 a.a.) | PeproTech | Cat# 100-18B-100UG |
| ReLeSR | Stemcell Technologies | Cat# 100-0483 |
| Rotenone, ≥95% | Sigma-Aldrich | Cat# R8875-1G |
| SB 431542 | Tocris | Cat# 1614 |
| Seahorse XF DMEM Medium pH 7.4 | Agilent Technologies | Cat# 103575-100 |
| Sodium Pyruvate (100 mM) | Thermo Fisher Scientific | Cat# 11360070 |
| Sucrose, for molecular biology, ≥99.5% | Sigma-Aldrich | Cat# S0389-500G |
| SuperSignal West Pico PLUS Chemiluminescent Substrate | Thermo Fisher Scientific | Cat# 34580 |
| Triton X-100 | Sigma-Aldrich | Cat# T8787 |
| Tryple Express Enzyme (1X), Phenol Red | Thermo Fisher Scientific | Cat# 12-605-010 |
| Tween® 20 | Sigma-Aldrich | Cat# P2287-500ML |
| XAV 939 | Tocris | Cat# 3748 |
| Y-27632 dihydrochloride | Tocris | Cat# 1254 |
| **Critical commercial assays** |  |  |
| STEMdiff™ Cerebral Organoid Kit | STEMCELL Technologies | Cat# 8570 |
| DNeasy Blood & Tissue Kit (250) | Qiagen | Cat# 69506 |
| PyroMark PCR Kit (200) | Qiagen | Cat# 978703 |
| PyroMark Q48 Advanced Reagents (4 x 48) | Qiagen | Cat# 974002 |
| PyroMark Q48 Magnetic Beads (300) | Qiagen | Cat# 974203 |
| Seahorse XFe96/XF Pro FluxPak Mini | Agilent Technologies | Cat# 103793-100 |
| TUNEL Assay Kit (Fluorescence, 488 nm) | Cell Signaling Technology | Cat# 25879S |
| **Critical commercial assays** |  |  |
| SCrnaSEQ DATA |  |  |
| **Experimental models: cell lines** |  |  |
| ipsc P1-12% het | Chichagova et al. | N/A |
| ipsc P1-58% het | Chichagova et al. | N/A |
| ipsc P1-72% het | Chichagova et al. | N/A |
| ipsc P2-0% het | Gunnewiek et al. | N/A |
| ipsc P2-15% het | Gunnewiek et al. | N/A |
| ipsc P2-42% het | Gunnewiek et al. | N/A |
| ipsc P2-64% het | Gunnewiek et al. | N/A |
| **Biological samples** |  |  |
| Paraffin-embedded tissue from healthy and m.3243A>G brain of human origin |  |  |
| **Software and algorithms** |  |  |
| QuantaSoft™ Software | Bio RAD | RRID:SCR_025696 |
| Seahorse Wave | Agilent Technologies | RRID:SCR_014526 |
| PyroMark Q24 Software v2.0.8 | Qiagen | N/A |
| Imaris 9.3 | Oxford Instruments | RRID:SCR_007370 |
| Affinity photo 2 | Affinity | RRID:SCR_016951 |
| Adobe Illustrator 2024 | Adobe | RRID:SCR_010279 |
| QuPath 0.5.1 | https://qupath.github.io/ | RRID:SCR_018257 |
| FIJI |  | RRID:SCR_002285 |
| CellProfiler 4.2.8 | https://cellprofiler.org | RRID:SCR_007358 |
| Cell Ranger |  | RRID:SCR_017344 |
| R studio |  | RRID:SCR_000432 |
| GraphPad Prism 10 |  | RRID:SCR_002798 |
| **Other** |  |  |
| NuPAGE™ Novex™ 4–12% Bis-Tris Protein Gels | Thermo Fisher Scientific | Cat# NP0335BOX, NP0336BOX |
| NuPAGE™ MES SDS Running Buffer | Thermo Fisher Scientific | Cat# NP0002 |
| iBlot™ 2 Transfer Stacks, PVDF, mini | Thermo Fisher Scientific | Cat# IB24002 |
| Ibidi USA µ-Dish 35 mm | Ibidi | Cat# 80136 |
| ddPCR Supermix for Probes (no dUTP) | Bio-Rad | Cat# 1863023 |
| Automated droplet-generating oils | Bio-Rad | Cat# 1864110, 1863004 |
| DG32 automated droplet generator cartridges | Bio-Rad | Cat# 1864108 |

### LEAD CONTACT AND MATERIALS AVAILABILITY

Requests and detailed information of reagents will be made available by the Lead Contacts, Denisa Hathazi and Rita Horvath.

### EXPERIMENTAL MODEL AND SUBJECT DETAILS

#### Ethical approval

We use human primary fibroblast derived induced pluripotent stem cells (iPSCs) to generate cortical organoids and neuronal cells. Human brain autopsy of a patient with MELAS syndrome and age and sex match controls was also studied. All patients have been consented for using their clinical data and biological samples for this research as part of the “Genotype and phenotype in inherited neurodegenerative diseases” study (REC ID: 13/YH/0310, Yorkshire & Humber – Bradford Leeds Research Ethics Committee).

The biological samples have been securely stored in approved research facilities at the University of Cambridge, or within the Neuropathology Laboratory of the Cambridge University Hospitals NHS Foundation Trust.

#### iPSC culture

Human induced pluripotent stem cells (iPSC) were a generous gift from Valeria Chichagova, David Steel and Majlinda Lako, Newcastle University, Newcastle, UK (Patient 1 clones 12%, 58% and 72%)^1^ and Tamas Kozicz and Eva Morava-Kozicz from Icahn School of Medicine at Mount Sinai, New York, US (Patient 2 clones 0%, 15%, 42% and 64%)^2^ with additional details in Table S1. All cell lines were adapted and cultured in mTeSR on plates coated with Geltrex. Media was changed daily and colonies were passaged using ReLeSR according to manufacturer’s protocol when confluency reached 60-70%. For cryopreservation cells were frozen in 90% mTeSR Plus containing 10% dimethyl sulfoxide-DMSO.

#### Generation of cerebral organoids and air liquid interface slices (ALI-COs)

For the generation of cortical organoids we have employed a method previously published^3,4^ utilising the STEMdiff Cerebral Organoid kit. Briefly, iPSCs were detached using Accutase at 37°C for 5 minutes and then washed using mTeSR and 18,000 cells were plated in bottom rounded dishes in Media 1 +Supplement A. Media change was performed at day 3. At day 5 embryoid bodies (EBs) were moved in Media 1+Supplement B which supports the formation of neuroectoderm. At day 7 EBs were embedded in 20 µL of Matrigel and put in Media2+ Supplement C and D until day 10. From day 10 onwards organoids were moved to Media 2+Supplement E and organoids kept on an orbital shaker. Media was changed every 4 days and from 35 DIV onwards, Matrigel was added in 1:50 dilution to achieve a polarized cortical plate formation. Between 45-50 DIV 300 µm ALI-COs were prepared using a vibratome and placed on Millicell-Cell culture inserts, 30 mm, inserts. After slicing, ALI-COs were fed daily using a Slice Media containing Neurobasal medium supplemented with 1× B27 supplement, 0.45% (w/v) glucose (Sigma-Aldrich, G8769), 1× Glutamax and 1% antibiotic– antimycotic.

#### Organoid dissociation into single cells

Organoid cell dissociation was performed using both mechanical and enzymatic methods similar to previously described^5^. Organoids were transferred into a dish containing 1xdPBS. Washed slices were transferred into a gentleMACS C tube containing 2 ml of papain solution (20 units per ml; Worthington, PAP2) and ran on a gentleMACS Octo dissociator (Miltenyi) using the default ABDK program. After dissociation the cell suspension was diluted with dPBS containing 0.5LmgLml^−1^ DNAse and then spun down at 300L×L*g* for 5 min. The cell pellet was resuspended and filtered using a 70 μm strainer and centrifuged again. Resulted cell suspension was centrifuged using the above described conditions and then resuspended in the appropriate buffer. For scRNA seq the cell pellet was resuspended in dPBS (445 cells per µl) containing 0.04% BSA. For the Seahorse experiment cells were resuspended in N2B27 media containing 10 μm Y-27632 and plated on Seahorse plates (100,000 cells/well) pre-coated with polyethylenimine-PEI and Geltrex.

#### FACS sorting of neurons and astrocytes from organoids

For cell sorting, organoids were washed in 1x dPBS, before being placed into gentleMACSTM C tubes in 2 ml of TrypLE for tissue dissociation. Dissociated single cells were then stained with a cell viability dye (Zombie-NIR) and labelled with astroglia (HEPACAM) or neuron (L1CAM) specific antibodies before FACS using the following protocol. Cells were resuspended in dPBS containing primary antibodies diluted in 0.05% BSA and samples were incubated for 1 hour on a shaker at 37 °C. The samples were then washed in 0.5% BSA dPBS and centrifuged for 5 minutes at 300x g, before resuspended in the secondary antibody solution and incubated for another hour on the shaker at 37 °C. After the final wash, cells were resuspended at 200 cells ml−1 and sorted using a BD FACSAria™ Flow Cytometer. Automatic normalization was applied using individually stained samples and a negative control. The gating was determined using an isotype control (mouse IgG1 antibody). Cells were filtered for doublets and debris, and only HEPACAM positive astroglia or L1CAM positive neurons were retained. Post-sorting, samples were centrifuged at 2000x g for 1 minute at 4 °C, followed by flash freezing on dry ice and subsequent storage at -80 °C for long-term preservation.

#### Seahorse assay

We measured the oxygen consumption rate (OCR) using the Seahorse XFe96 Analyzer following manufacturer’s instructions. Three slices from different organoids were dissociated as described above and then 100,000 cells/ well were seeded in a Seahorse 96-well plate (minimum 20 wells per group). Cell were left for 3-5 days to recover after dissociation in N2B27 media. Prior to measuring cultures were switched to Seahorse XF DMEM Medium pH 7.4, supplemented with 5 mM Glucose, 1 mM sodium pyruvate, 2 mM L-Glutamine. Cells were maintained at 37C in a CO_2_-free incubator for one hour prior to OCR measurement. We measured simultaneously mitochondrial respiration (oxygen consumption rate, OCR) and anaerobic glycolysis (extracellular acidification rate, ECAR) using the sequential introduction of oligomycin, FCCP, and then rotenone and antimycin (1 µM for all). The addition of these drugs allowed us to calculate: basal OCR level, basal ECAR level, maximal respiration, and ATP production, as described in detail before. Data was normalized to total DNA (Hoechst 33342, 2 μM) and analysed using WAVE software (Agilent Technologies).

#### Generation of 2D cortical neurons

Neuronal induction was performed based on a protocol described by Maroof et all..^6^ When iPSC have reached 70% confluency cells were detached using Accutase as described above and then plated in a 6 well pate pre-coated with Geltrex. Cells were kept in mTeSR supplemented with 10Lμm Y-27632 for 24 hours after which the media was switched with neural induction media NI containing N2B27 media supplemented with 2µM XAV939, 100nm LDN193189 and 10µM SB431542. Daily media changes were performed and after 10 days were detached using Accutase (as described above) and replated on a T25 flask coated with Geltrex in N2B27 media. After 5 days NPCs were replated (using Accutase) in 6 well plates Geltrex pre-coated plates and maintained in N2B27 supplemented with 100nm FGF2. For neuronal differentiation NPCs were plated at 50,000 cells/cm^2^ on PEI and Geltrex pre-coated plates in N2B27 media supplemented with 10μm Y-27632. 24 hours after plating cell media was replaced with N2B27 containing 10µm Compound E which induced cell differentiation. Cell media was replaced every 3 days and cells were kept in Compound E for 7 days. Afterwards cells were slowly transitioned from N2B27 to BrainPhys supplemented with 1X SM1 without vitamin A, 1X N-2 Supplement and 10 ng/ml BDNF. Cell were matured for a total of 30 days.

#### Cell viability

Cell viability was measured using the PrestoBlue assay, following the manufacturer’s recommendation. 2D neurons were seeded in 96-well plates and washed with BrainPhys medium before adding 200 µL of BrainPhys containing 5% PrestoBlue solution per well. The plates were then incubated at 37°C with 5% COL for 4 hours. After incubation, the PrestoBlue/media solution was carefully transferred to a new 96-well plate for measurement, while the remaining cells were replenished with 200 µL of fresh BrainPhys medium and returned to the incubator under the same conditions for an additional 4 hours. Fluorescence readings were obtained using the BMG FLUOstar Omega Microplate Reader (BMG Labtech) according to the manufacturer’s specifications. Blank correction was performed using BrainPhys medium containing 5% PrestoBlue incubated without cells, and values were recorded for further analysis.

For metabolic stress conditions, the same cells were treated with 10 µM GSK-2837808A and 0.1% AlbuMAX in BrainPhys medium for 12 hours before performing the PrestoBlue assay. Fluorescence measurements were taken as described for the basal conditions.

#### m.3243A>G mtDNA heteroplasmy measurements

DNA was extracted from whole organoid slices using the DNeasy Blood and Tissue Kit from according to the manufacurer’s protocol and quantified using a NanoDrop spectrophotometer (Thermos Scientific). Heteroplasmy measurements were performed using the QIAGEN Q48 Autoprep workstation system. A section of the mitochondrial genome containing the mutation of interest (m.3243A>G) was amplified from 50 to 100 ng extracted DNA using the PyroMark PCR Kit. Primers were designed according to manufacturer’s specifications and sequences are found in Table S2. After confirmation of successful amplification by gel electrophoresis, heteroplasmy levels were analysed by running the PCR product on Q48 Autoprep workstation along with the appropriate sequencing primer and following the manufacturer’s protocol. Heteroplasmy was then called using the Pyro Mark analysis software and reported as the percentage of mutant base present in the samples.

A PCR-based assay was also used to determine the proportion of wild-type to mutant mtDNA. Specific primers were used to amplify the region where the mtDNA mutation is located and the dinucleotide substitution in the forward primer creates a restriction site for the restriction enzyme ApaI, which can cleave the mutant sequence (GGGCCC) when the m.3243A>G mutation is present. PCR amplification was carried out on a BioRad PCR Thermal Cycler using the following protocol: 1 min denaturation at 94°C, 1 min annealing at 55°C, and 45 s extension at 72°C for 25Lcycles. To confirm the presence of the mtDNA mutation, 237 bp PCR samples were made, and the unilateral sample was digested with ApaI at 37°C for 60 min. These samples were run on a 2% Agarose gel and visualized using GelRed nucleic acid stain using. ApaI digestion produces two units, one of 127 bp and one of 110 bp, which can easily be differentiated from the uncut wildLtype unit by gel electrophoresis.

#### Quantification of mitochondrial DNA copy numbers in whole organoids using qPCR

The relative mtDNA copy number per cell was quantified by a multiplex Taqman qPCR assay by amplifying *MT-ND1* (mitochondrial encoded gene) and *B2M* (nuclear encoded gene) with a CFX96™ Real-Time PCR Detection System, Bio-Rad following the protocol described previously^7^. The primers used for template generation of standard curves and the qPCR reaction are described in Table S2. The copies per µL of each template were standardized to 1 * 10^10^ and a serial dilution in 1Log_10_ dilution steps, was amplified along with the DNA negative control on each qPCR plate. This was performed in 20 µL reactions in a 96 well-plate, sealed using microplate ‘B’ plate sealers. The reaction mixture was composed of: 5 µL x 5 x Taqman (Bio-rad 4369510), 0.4 µM of reverse and forward primers, 25-50 ng of DNA template, 0.2 µL MyTaq HS DNA polymerase and PCR-grade autoclaved sterile deionised water (to make up 20 µL reaction mixture). The cycling conditions were as follows: 1) initial denaturation at 95°C for 3 minutes, 2) 40 cycles of denaturation at 95°C for 10 seconds, 3) annealing and extension at 62.5°C for 1 minute. The relative mtDNA copy number was calculated using the ΔCt data following the equation: CopyNumber = 2 (2^-ΔCt^) where Delta Ct (ΔCt) equals the sample Ct of the mitochondrial gene (MTND1) subtracted from the sample Ct of the nuclear reference gene (B2M).

#### Quantification of mitochondrial DNA copy numbers in neurons and astrocytes using ddPCR

The determine the mtDNA absolute copy numbers by digital droplet PCR we have used the protocol described by Burr et al.^8^ Following Proteinase K treatment, samples were diluted with ddH_2_O 1:125 to obtain the final concentration of 10 cell equivalents per μl. 1 μL cell lysate was combined with locus-specific primers, FAM-labelled MT-ND4 (mtDNA) and HEX-labelled RNAseP (nuclear DNA) probes (Table S2), and digital droplet PCR Supermix for probes (no dUTP). Cycling conditions for the probes were tested to ensure optimal annealing/extension temperature and optimal separation of positive from empty droplets. Optimization was done with a known positive control. All reactions were performed on a QX200 ddPCR system, and each sample was evaluated in technical duplicates or triplicates. Reactions were partitioned into a median of ∼10,000 droplets per well using the QX200 droplet generator. Emulsified PCRs were run on a 96-well thermal cycler using cycling conditions identified during the optimization step (95°C 10 seconds; 40 cycles of 94°C 30 seconds and 54, 56, or 60°C 1 second; 98°C 10 seconds; 4°C hold). Plates were read and analysed with the QuantaSoft software to assess the number of positive droplets for the gene of interest, reference gene, or neither. mtDNA copy numbers per cell were calculated as follows: MT-ND4 droplet number / RANaseP ^∗^ 2.

#### Protein lysate preparation and immunoblotting analysis

Whole organoid slices were lysed in 200 µL of lysis buffer (pH 7.8) 150 mM NaCl, 1 % SDS, and Complete Mini-Roche using a plastic douncer. Then samples were centrifuged for 5 min at 4°C and 5000 *g*. Protein concentration of the supernatant was determined by BCA assay (according to the manufacturer’s protocol).

For immunoblot studies, 10 μg protein was used in each case, loaded on a gradient polyacrylamide gel NuPage 4-12% Bis-Tris Protein gels, and separated for 120 min at 120 V. Following the separation, proteins were transferred to PVDF membrane using the iBlot2 system from ThermoFisher according to the manufactures protocol. Membranes were blocked with 5% Milk prepared in PBS-T for 2 h followed by four washing steps using PBS-T with 0.1% Tween20 (PBS-T). Membranes were incubated with several primary antibodies (Table 1) at 4 °C (overnight) and then washed in PBS-T thrice. Horseradish peroxidase conjugated secondary goat anti-rabbit antibody or goat anti-mouse antibody was diluted at 1:25,000 and added to membranes for 1 h. Next, membranes were washed three times in PBS-T for 10 min. By using the enhanced chemiluminiscence, horseradish peroxidase substrate (Super-Signal West Pico and Amersham ECL Select Western Blotting Detection Reagent) signals were detected using a Amersham 680.

#### Immunocytochemistry

Organoid slices and whole organoids at 30 and 75 days in vitro (DIV) were fixed in 4% paraformaldehyde (PFA) at room temperature under gentle agitation for 45, 25, and 30 minutes, respectively. Following fixation, samples were washed three times with phosphate-buffered saline (PBS) and incubated in 30% sucrose in PBS for at least 24 hours or until they sank. Fixed samples were embedded in Optimal Cutting Temperature (OCT) compound and processed into 12-µm frozen sections for immunostaining.

Sections were encircled with a hydrophobic barrier (Pap Pen), washed in PBS (5 minutes), and permeabilized with 0.5% Triton X-100 for 10 minutes. Blocking was performed at room temperature for 1 hour in 10% goat or donkey serum in PBS, depending on the species of the secondary antibody. Sections were incubated overnight at 4°C with primary antibodies diluted in 5% goat or donkey serum in PBS. The following day, sections were washed in PBS and incubated with the corresponding secondary antibody (5% goat or donkey serum in PBS) for 2 hours at room temperature. Nuclei were counterstained with 4,6-diamidino-2-phenylindole (DAPI; 0.2 mg/ml), and samples were mounted using ProLong Gold Antifade Mountant.

Apoptosis staining was performed using the TUNEL assay according to the manufacture’s protocol. After fixation samples rinsed two times with PBS followed by permeabilization in PBS-TB (0.2% Triton X-100, 0.5% BSA in PBS) for 30 minutes at room temperature. Samples were wash twice with PBS and incubated TUNEL equilibration buffer followed by incubation with TUNEL reaction buffer and TdT Enzyme for 1 hour at 37°C. After this samples were co-stained with primary and secondary antibodies as described above.

2D cell cultures were fixed in 4% PFA for 10 minutes at room temperature, followed by permeabilization with 0.4% Triton X-100 in PBS for 10 minutes. Blocking was performed for 1 hour in 5% bovine serum albumin (BSA) in PBS. Cells were then incubated overnight at 4°C with primary antibodies diluted in 1% BSA in PBS, followed by secondary antibody incubation for 1 hour at room temperature. Samples were mounted using ProLong Gold Antifade Mountant with DAPI.

Formalin-fixed, paraffin-embedded (FFPE) 10-µm human brain tissue sections, provided by Dr. Kieren Allison, were first baked at 60°C for 2 hours to remove paraffin. Sections were then deparaffinized in xylene for 45 minutes, followed by sequential washes in 100%, 96%, 70%, and 50% ethanol (10 minutes each). After rehydration in distilled water, antigen retrieval was performed using 10 mM sodium citrate buffer (0.05% Tween-20, pH 6.0) at 90°C for 20 minutes. Following antigen retrieval, sections were washed three times in wash buffer (PBS + 0.3% Triton X-100) and blocked for 2 hours in 5% BSA in wash buffer. Samples were incubated overnight at 4°C with primary antibodies diluted in 5% BSA in PBS. The next day, sections were washed twice in wash buffer, followed by two PBS washes, and incubated with Lipofuscin Autofluorescence Quencher for 1 minute, followed by additional PBS washes. Finally, sections were incubated with secondary antibodies in PBS, washed, and mounted using ProLong Gold Antifade Mountant with DAPI.

Information about antibody dilutions can be found in Table S3.

#### Image acquisition and processing

Stained organoid microsections and cells were imaged using a 20×, 40×, or 60× objective on a Nikon Eclipse Ti-E inverted microscope, equipped with a 4-channel Integrated Laser Engine (ILE-400) and an Andor Dragonfly 200 spinning disk system. The system included two iXon Ultra 888 EMCCD cameras and was operated using Fusion software. Images were acquired at Image frame size was 1024 × 1024 pixels, with automatic Z-step adjustments based on tissue thickness. Whole organoid microsections were imaged using an Opera Phenix Plus High content Screening System from Revvity, using a 40 x water objective. For all experiments the confocal optical mode was used and the autofocus was set to two peaks. The binning was set to two and the camera ROI to 1024 x 1024. A total of 20 stacks were taken with 2 µm distance for each stack. Mitochondrial morphology analysis was conducted using a Zeiss LSM-980 confocal microscope equipped with an Airyscan2 detector to achieve super-resolution imaging. Images were acquired in super-resolution mode, with a bit-depth of 16 bits, covering regions of 28 × 28 µm for cell bodies and 130 × 12 µm for axons.

To ensure complete coverage of cellular structures, Z-stack acquisition was performed with 5 slices at 400 nm intervals for cell bodies and 4 slices at 300 nm intervals for axons. All images were obtained using frame scanning mode. Camera exposure and gain were kept identical while collecting images for each experiment. For illustration purposes, the recommended guidelines were followed and images were only minimally processed in Imaris or Affinity Photo 2 (only brightness and/or contrast) without affecting data presentation. For each sample all channels were uniformly adjusted.

For the neuronal layering analysis of CTIP2 and SATB2 cells in whole organoid microsections post-acquisition image analysis was performed using StarDist^9^ (https://github.com/stardist/stardist) in QuPath^10^ (0.5.12.; https://qupath.github.io/). Raw imaging data acquired from the Opera Phenix high-content screening system was exported from Harmony software, then Z-projected and stitched at full resolution using Fiji (ImageJ) with the ‘Operetta Importer’ and the ‘Grid/Collection Stitching’ plugin. Images were imported as maximum intensity projections with a down-sampling factor of 1 to preserve high resolution. Illumination correction for the DAPI channel was applied in Fiji, setting the central intensity to 1 with increasing values toward the periphery to allow for a better nuclear segmentation. Nuclei segmentation was conducted in QuPath using a pre-trained StarDist model optimized for whole-slide image analysis (https://github.com/tpecot/WholeSlideImageAnalysisWithQuPath) ^11–13^. The QuPath StarDist extension was employed to detect nuclear boundaries based on DAPI staining, with segmentation parameters adjusted to a detection threshold of 0.5, a pixel size of 0.5 µm, and a minimum nucleus area of 30 µm² to exclude debris. Following segmentation, nuclei were evaluated for CTIP2 and SATB2 expression based on manually set fluorescence intensity thresholds, which were kept constant across all images. The percentage of CTIP2-positive or SATB2-positive nuclei relative to the total DAPI-positive nuclei was then calculated.

For mitochondrial morphology post-acquisition image analysis was carried out using Fiji (ImageJ) software, where background subtraction with a 50-pixel radius, despeckle filtering, and Gaussian blur with a 1-pixel radius were applied to enhance image quality. Mitochondrial segmentation within the cell body and axonal regions was performed based on TOM20 intensity thresholding, and the resulting binary masks were subjected to particle analysis to extract quantitative morphological parameters. These parameters included mitochondrial area, which measured the total mitochondrial coverage within the analysed regions; perimeter, representing the total boundary length of individual mitochondria to assess shape complexity; form factor, a measure of mitochondrial shape complexity calculated as the ratio of perimeter squared to area, with values approaching 1 indicating a more circular shape and lower values suggesting elongated mitochondria; and aspect ratio, defined as the ratio of the major to minor axis of mitochondria, providing insights into their elongation and branching behaviour. Additionally, mitochondrial density was calculated as the number of mitochondria per micrometer of axonal length to provide a quantitative assessment of mitochondrial distribution along the axon.

Axonal swellings were quantified using CellProfiller (version 4.2.8; https://cellprofiler.org) with a custom image analysis pipeline. To enhance the visibility of the swellings, images were first pre-processed using the Smooth module with the Smooth Keeping Edges method, applying an edge intensity difference threshold of 0.05. The swellings were then identified using the IdentifyPrimaryObjects module, with a diameter range set between 2-25 pixels. An adaptive two-class Otsu thresholding method was applied. The objects were subsequently analysed using the MeasureObjectSizeShape module to extract morphological measurements. Finally, identified objects were filtered using the FilterObjects module, retaining those with a form factor between 0.7 and 1 and an area between 6-170 pixels to exclude artifacts and background noise. Thee areas of interest from two different batches were analysed for each sample.

#### scRNA sequencing pipeline and analysis

Reads were aligned to the GRCh38 human genome using Cell Ranger (version 7.0.1) that additionally performs filtering, barcode and UMI counting to obtain barcode and feature matrices. CellRanger detected 88.4% fraction reads on average per cell (ranging from 83.7% to 92.0%) and 1443-4368 (3383 on average) median genes per cell. Bias from false cell discovery due to potential ambient RNA contamination/low UMI counts was not eliminated in any of the samples. A conda environment has been created to install all required software, among which R and Satija’s Seurat R package-version 4.0/5.0.0^14^, R version 4.2.2.

To remove low quality cells and ensure high-quality data, cells with more than 20% of mitochondrial or ribosomal reads have been removed; the ratio between the log (number of genes) and log (number of molecules) has been evaluated for each cell; only cells that have a high score of such value (>0.8) and a high number of detected molecules (>1000) and a detected number of genes between 800 and 10000 have been kept. To remove low detected genes, those that are expressed in less than 10 cells have been removed. Additionally, long non-coding genes, antisense transcripts and pseudogenes have been removed from the analysis. The resulting dataset has 46725 genes. Normalisation and integration have been performed with sctranform, an R package pipeline, using the top 3000 features as identified by the SelectIntegrationFeatures function, to perform the integration step. To reduce the dimensionality of the dataset, a PCA and Uniform Manifold Approximation and Projection (UMAP) have been performed, the latter using the first 20 dimensions of the dataset, as determined with the Elbow method. The cell cycle stage has been determined using the CellCycleScoring function of the Seurat package.

To identify the cell clusters, we have employed the FindClusters method, selecting the Louvain algorithm with multi-level refinement, setting the clustering resolution to 0.185. This led to the identification on one cluster of stressed cells that was removed from further analysis. The final size of the dataset is of 119.591 cells. Clustering has been performed a second time, setting the clustering parameter to 0.18. To annotate the cell types we calculated the top 50 marker genes per cluster, using the FindAllMarkers Seurat function, using the Wilcoxon test and selecting only the positively expressed genes with a logarithmic fold change greater than 0.6 and a minimum percentage of cells in each cluster that express the gene of 0.6.

Cell maturity analysis was achieved by independently projecting data from our 100DIV organoids onto one scRNA-seq data set from fetal brain^15^ using scmap (v1.20.2) software^16^. We employed scmapCluster function of the scmap R package (‘scmapCluster()’ thresholdL□=□0)).

Monocle3 (v1.3.1) was employed for trajectory reconstruction, leaving the default parameters for the order_cells function and choosing RG as starting cell type.

Transcription factory activity analysis was performed using pyscenic (v0.12.0), python version 3.8.13, following the standard pipeline (GRNboost2, AUCell), leaving the default parameters values.

The Wilcoxon test was used to compare distribution of continuous variables between groups, for example gene expression. The Pearson’s chi-squared test was used to compare distributions of categorical data. P-values have been adjusted using the Benjamini-Hochberg procedure.

To identify genes that change in expression across conditions, we performed a differential expression analysis. Genes were identified with the FindMarkers function, setting the minimum number of cells expressing the gene to 3. Genes are considered differentially expressed (DEG) if the adjusted p-values is smaller than 0.05 and the logarithmic fold change is greater or smaller than 0.5. Overlap of the significant terms with the Mitocarta database (version 3.0) has been performed.

Pathway enrichment analysis was performed using ClusterProfiler (v4.6.2) on gene sets identified by the differential expression analysis. Following are the parameters set for the enrichGo function: org.Hs.eg.db (v3.16.0) as reference organism database, p-values have been corrected with the “BH” method, p-value and q-value cutoffs are set to 0.05, and the “MF”, “BP”, “CC” ontologies have been selected.

#### Statistics

Sample sizes using iPSC lines and organoids were estimated from previous studies^17,19^. For this study we derived 5 independent batches of organoids from each iPSC line. For our experiments organoids used are from different batches unless specified. For some experiments such as heteroplasmy measurements we wanted to determine iter and intra batch variation. Experiments were repeated three times (or two times absolute quantification of mtDNA in sorted neurons and mitochondrial morphology in 2D neurons), which included at least three independent biological replicates, and all had similar results. At least three independent biological replicates were used per group for all statistical analyses in biological experiments. For Seahorse experiments, all assays were performed three times to confirm the reproducibility of the observed patterns. The reported n = 5 refers to five independent organoids run on the same plate to minimize inter-plate variability, which is a standard practice for Seahorse assays.

Errors bars displayed on graphs represent the mean +/− standard error of the mean (S.E.M.). Number of experiments and cells analyzed is shown in the corresponding paragraph in material and methods. Statistical significance was analyzed using the tests described in the corresponding figure legends. All statistical analyses and super-plots from the experimental data were performed using GraphPad (version 10) while scRNA-seq were performed using R. When normality was not assumed or defined, non-parametric tests were used. Unless stated otherwise, statistical significance was accepted at *P*LJ<LJ0.05, and the exact *P* values are included in the graphs. Where no significant differences were detected among groups, the overall ANOVA p-value is reported, and non-significant values are not shown in the plot.

##### Data S1 - Source Data

Uncropped western blotting images used for Figures 1, 3 and 6.

## SUPPLEMENTARY FIGURES

**Figure S1. A.** Representative agarose gel for determining the m.3243A>G mutation in iPSCs. Screening of the m.3243A>G mutation was done by PCRLJRFLP where a 528 bp PCR fragment is digested with ApaI. The wildLJtype PCR product does not contain ApaI restriction site, whereas PCR product containing the m.3243A>G mutation is cleaved into two fragments of 234 and 294 bp in length; **B.** Phase contrast imaging of whole organoids where expanded neuroepithelia show similar bud-formation between cortical organoids at 10 DIV; **C.** Representative immunofluorescence images of cortical organoids at DIV 30, showing the palisade-like organization of SOX2LJ progenitor cells (violet), surrounded by immature TUJLJ neurons (red) and NESTINLJ progenitor cells (green). Scale bar = 50 µm; **D.** Representative immunofluorescence images of ALI-COs at DIV 100, showing TUJLJ (green) neurons and GFAPLJ (red) glia across three biological replicates per group. Scale bar = 200 µm;

**Figure S2. A.** Representative immunofluorescence images of cortical plate regions at DIV 75, showing the forebrain identity marker FOXG1 (green) and early-born CTIP2LJ neurons (violet), with insets highlighting cortical plate organization. Scale bar = 50 µm; **B.** Average gene expression of marker genes for various subpopulations identified in 11 unbiased clusters by scRNA-seq within the merged dataset of control and C9 ALI-COs (nLJ=LJ119,591 cells). Cell annotation: ULN, upper-layer cortical neurons; DLN5B and DLN5A, deep-layer neurons (layer 5, subtypes A and B); DLN6, deep-layer neurons (layer 6); IN, interneurons; yULN, young upper-layer neurons; IP, intermediate progenitors; RG, radial glia; oRG, outer radial glia; CP, choroid plexus; Ui, unidentified cells; **C.** UMAP representations of scaled expression levels and distribution of key marker genes of major cell-types identified; **D.** Representative immunofluorescence images of laminar organization in cortical organoid slices at DIV 100, where SATB2LJ cells (red) mark ULNs and CTIP2LJ cells (green) label DLN5 neurons with scale bar= 200 µm; **E.** Bar plots indicate proportion of cells mapping to transcriptional profiles of various fetal ages in scMAP; F. UMAP representation of inferred developmental trajectories in low, medium and high heteroplasmy obtained by using Monocle3.

**Figure S3. A.** Quantification of WB data for SDHB (Complex II) and ATP5A (Complex V). Data were normalized to TOM20 to account for mitochondrial content. Data represent the mean ± SEM; *n* = 3 ALI-COs from three independent batches, with two organoids pooled per sample. Statistical analysis: one-way ANOVA with Tukey’s post hoc test (overall ANOVA *p*-values indicated in the graphs); **B**. Extracellular acidification rate measurements in 100 DIV dissociated organoids. The upper panel represents data from Patient 1, and the lower panel represents data from Patient 2. OCR was measured at basal levels, followed by supplementation with 1 µM oligomycin, 1 µM FCCP, and 1 µM rotenone/antimycin A; **C.** Quantification of mitochondrial respiration. Bar plots show Maximal respiration, proton leak and spear respiratory capacity. The left panel represents data from Patient 1, and the right panel represents data from Patient 2. *n* = 5 dissociated ALI-COs for heteroplasmic and control organoids, with each experiment performed in triplicate to confirm trends. Statistical analysis: one-way ANOVA with Tukey’s post hoc test (overall ANOVA *p*-values indicated in the graphs); **D.** Cytometric analysis of L1CAM immunoreactivity for neurons (left pannel) and of HEPACAM for astroglia (right panel). Graphs show also gating strategy (red and blue); **E.** Box plots showing absolute quantification of mtDNA in astrocytes sorted from 100 DIV organoids. *n* = 2 independent sorting experiments from three organoids pooled per sample. Statistical analysis: one-way ANOVA with Tukey’s post hoc test (overall ANOVA *p*-values indicated in the graphs); F. Heteroplasmy levels of sorted neurons and astrocytes determined by pyrosequencing.. *n* = 2 independent sorting experiments from three organoids pooled per sample.

**Figure S4. A.** Venn diagram illustrating shared and unique transcription factors (TFs) identified in high, medium, and low heteroplasmy organoid groups. TF analysis was performed using SCENIC on merged data, representing transcription factors across all cell populations; Purple square emphasizes the common TFs identified in all heteroplasmic samples; **B.** Dot plots of all identified TFs in DLN5B in high, medium and low heteroplasmy; **C.** Venn diagram illustrating shared and unique transcripts for low and high DLN5B and ULN; **D.** Box plots with individual data points representing GO enrichment of major pathways in DLN5B vs. ULN for both low and high heteroplasmy. The Wilcoxon test was used to compare the distribution of continuous variables between groups, such as gene expression levels.

**Figure S5. A.** Skeletal muscle histopathology. H&E and Gomori trichrome staining (upper panels) of transverse muscle sections from a MELAS patient reveal fiber-size heterogeneity, fibrosis, rods, and fibers with multiple internalized nuclei and cores. The identification of muscle fiber type in a cross-section of a muscle biopsy from a MELAS patient was performed using immunohistochemical analysis (lower panel). Muscle fiber types were distinguished by myosin ATPase staining at different pH levels using peroxidase immunohistochemistry, while oxidative muscle fibers were identified by succinate dehydrogenase (SDH) activity; **B.** Brain histopathology of a MELAS patient. H&E staining in the occipital lobe (upper panel) and cortex (lower panel) shows severe neuronal loss. Representative GFAP immunohistochemistry staining in the human brain of a MELAS patient reveals intense astrogliosis, characterized by widespread reactive astrocytes with strong GFAP expression, indicative of a neuroinflammatory response. Scale bar: 100 µm; **C.** Representative immunofluorescence images of neuronal progenitor cells (NPCs) derived from iPSCs. SOX2LJ cells (red) mark neural stem and progenitor cells, while NESTINLJ cells (green) label intermediate filament proteins. Scale bar: 50 µm; **D.** Bar plots quantifying the number of SOX2LJ and NESTINLJ cells, normalized to the total number of cells, indicating the presence of undifferentiated, proliferative NPCs. Data represent the mean ± SEM; *n* = 3 independent batches from three different neuronal inductions. Statistical analysis: one-way ANOVA with Tukey’s post hoc test (no significant difference observed); **E.** Representative immunofluorescence images of 2D cortical neurons showing that the majority of cells are positive for MAP2, a neuronal marker, and CTIP2, a deep-layer neuronal marker. *n* = 3; **F.** Bar chart showing relative cell viability determined by the PrestoBlue assay under basal and stressed conditions, where mitochondrial function was acutely stimulated using 10 µM GSK- 2837808A + 0.1% AlbuMAX for 12 hours. Data represent the mean ± SEM; n = 6 from three independent batches of neurons. Statistical analysis: two-way ANOVA with Tukey’s post hoc test for multiple comparisons (no significant difference observed); **G.** Representative immunofluorescence images of 2D control and high-heteroplasmy neurons. MAP2 (magenta) marks dendrites and cell bodies, while TOM20 (green) marks mitochondria. Scale bar: 50 µm.

## Supplemental tables

**Table S1.** Supplementary information on the subjects related to the iPSC lines used in this study.

**Table S2.** List of primers used, as referenced in the STAR Methods.

**Table S3.** List of antibodies and their dilutions used in this study, as referenced in the STAR Methods.

**Table S4.** Top 50 marker genes per cluster identified from the merged scRNA-seq dataset.

**Table S5.** Differentially expressed gene transcripts in deep-layer neurons (DLN5B) when comparing high versus low heteroplasmy conditions, based on scRNA-seq analysis.

**Table S6.** Differentially expressed genes in deep-layer neurons (DLN5B) when comparing high versus low heteroplasmy conditions, based on scRNA-seq analysis.

**Table S7.** Differentially expressed genes in high-heteroplasmy deep-layer 5B neurons compared to upper-layer neurons.

**Table S8.** Differentially expressed genes in control-heteroplasmy deep-layer 5B neurons compared to upper-layer neurons.

## REFERENCES

1. Chinnery, P.F. (1993). Primary Mitochondrial Disorders Overview. In GeneReviews((R)), M.P. Adam, J. Feldman, G.M. Mirzaa, R.A. Pagon, S.E. Wallace, and A. Amemiya, eds.

2. Parikh, S. (2010). The neurologic manifestations of mitochondrial disease. Dev Disabil Res Rev 16, 120–128. 10.1002/ddrr.110.

3. Dunn, D.A., Cannon, M.V., Irwin, M.H., and Pinkert, C.A. (2012). Animal models of human mitochondrial DNA mutations. Biochim Biophys Acta 1820, 601–607. 10.1016/j.bbagen.2011.08.005.

4. Olkhova, E.A., Smith, L.A., Bradshaw, C., Gorman, G.S., Erskine, D., and Ng, Y.S. (2023). Neurological Phenotypes in Mouse Models of Mitochondrial Disease and Relevance to Human Neuropathology. Int J Mol Sci 24. 10.3390/ijms24119698.

5. Keshavan, N., Minczuk, M., Viscomi, C., and Rahman, S. (2024). Gene therapy for mitochondrial disorders. J Inherit Metab Dis 47, 145–175. 10.1002/jimd.12699.

6. Hathazi, D., and Horvath, R. (2023). Mitochondrial DNA editing with mitoARCUS. Nat Metab 5, 2039–2040. 10.1038/s42255-023-00933-5.

7. Nash, P.A., and Minczuk, M. (2023). Manipulation of Murine Mitochondrial DNA Heteroplasmy with mtZFNs. Methods Mol Biol 2615, 329–344. 10.1007/978-1-0716-2922-2_23.

8. Shoop, W.K., Lape, J., Trum, M., Powell, A., Sevigny, E., Mischler, A., Bacman, S.R., Fontanesi, F., Smith, J., Jantz, D., et al. (2023). Efficient elimination of MELAS-associated m.3243G mutant mitochondrial DNA by an engineered mitoARCUS nuclease. Nat Metab 5, 2169–2183. 10.1038/s42255-023-00932-6.

9. Pickett, S.J., Grady, J.P., Ng, Y.S., Gorman, G.S., Schaefer, A.M., Wilson, I.J., Cordell, H.J., Turnbull, D.M., Taylor, R.W., and McFarland, R. (2018). Phenotypic heterogeneity in m.3243A>G mitochondrial disease: The role of nuclear factors. Ann Clin Transl Neurol 5, 333–345. 10.1002/acn3.532.

10. Ng, Y.S., Lax, N.Z., Blain, A.P., Erskine, D., Baker, M.R., Polvikoski, T., Thomas, R.H., Morris, C.M., Lai, M., Whittaker, R.G., et al. (2022). Forecasting stroke-like episodes and outcomes in mitochondrial disease. Brain 145, 542–554. 10.1093/brain/awab353.

11. Latchman, K., Saporta, M., and Moraes, C.T. (2023). Mitochondrial dysfunction characterized in human induced pluripotent stem cell disease models in MELAS syndrome: A brief summary. Mitochondrion 72, 102–105. 10.1016/j.mito.2023.08.003.

12. Klein Gunnewiek, T.M., Van Hugte, E.J.H., Frega, M., Guardia, G.S., Foreman, K., Panneman, D., Mossink, B., Linda, K., Keller, J.M., Schubert, D., et al. (2020). m.3243A > G-Induced Mitochondrial Dysfunction Impairs Human Neuronal Development and Reduces Neuronal Network Activity and Synchronicity. Cell Rep 31, 107538. 10.1016/j.celrep.2020.107538.

13. Hamalainen, R.H., Manninen, T., Koivumaki, H., Kislin, M., Otonkoski, T., and Suomalainen, A. (2013). Tissue- and cell-type-specific manifestations of heteroplasmic mtDNA 3243A>G mutation in human induced pluripotent stem cell-derived disease model. Proc Natl Acad Sci U S A 110, E3622–3630. 10.1073/pnas.1311660110.

14. Winanto, Khong, Z.J., Soh, B.S., Fan, Y., and Ng, S.Y. (2020). Organoid cultures of MELAS neural cells reveal hyperactive Notch signaling that impacts neurodevelopment. Cell Death Dis 11, 182. 10.1038/s41419-020-2383-6.

15. Tolle, I., Tiranti, V., and Prigione, A. (2023). Modeling mitochondrial DNA diseases: from base editing to pluripotent stem-cell-derived organoids. EMBO Rep 24, e55678. 10.15252/embr.202255678.

16. Eichmuller, O.L., and Knoblich, J.A. (2022). Human cerebral organoids - a new tool for clinical neurology research. Nat Rev Neurol 18, 661–680. 10.1038/s41582-022-00723-9.

17. Szebenyi, K., Wenger, L.M.D., Sun, Y., Dunn, A.W.E., Limegrover, C.A., Gibbons, G.M., Conci, E., Paulsen, O., Mierau, S.B., Balmus, G., and Lakatos, A. (2021). Human ALS/FTD brain organoid slice cultures display distinct early astrocyte and targetable neuronal pathology. Nat Neurosci 24, 1542–1554. 10.1038/s41593-021-00923-4.

18. Revah, O., Gore, F., Kelley, K.W., Andersen, J., Sakai, N., Chen, X., Li, M.Y., Birey, F., Yang, X., Saw, N.L., et al. (2022). Maturation and circuit integration of transplanted human cortical organoids. Nature 610, 319–326. 10.1038/s41586-022-05277-w.

19. Giandomenico, S.L., Mierau, S.B., Gibbons, G.M., Wenger, L.M.D., Masullo, L., Sit, T., Sutcliffe, M., Boulanger, J., Tripodi, M., Derivery, E., et al. (2019). Cerebral organoids at the air-liquid interface generate diverse nerve tracts with functional output. Nat Neurosci 22, 669–679. 10.1038/s41593-019-0350-2.

20. Trapnell, C., Cacchiarelli, D., Grimsby, J., Pokharel, P., Li, S., Morse, M., Lennon, N.J., Livak, K.J., Mikkelsen, T.S., and Rinn, J.L. (2014). The dynamics and regulators of cell fate decisions are revealed by pseudotemporal ordering of single cells. Nat Biotechnol 32, 381–386. 10.1038/nbt.2859.

21. Trevino, A.E., Muller, F., Andersen, J., Sundaram, L., Kathiria, A., Shcherbina, A., Farh, K., Chang, H.Y., Pasca, A.M., Kundaje, A., et al. (2021). Chromatin and gene-regulatory dynamics of the developing human cerebral cortex at single-cell resolution. Cell 184, 5053–5069 e5023. 10.1016/j.cell.2021.07.039.

22. Kiselev, V.Y., Yiu, A., and Hemberg, M. (2018). scmap: projection of single-cell RNA-seq data across data sets. Nat Methods 15, 359–362. 10.1038/nmeth.4644.

23. Bravo Gonzalez-Blas, C., De Winter, S., Hulselmans, G., Hecker, N., Matetovici, I., Christiaens, V., Poovathingal, S., Wouters, J., Aibar, S., and Aerts, S. (2023). SCENIC+: single-cell multiomic inference of enhancers and gene regulatory networks. Nat Methods 20, 1355–1367. 10.1038/s41592-023-01938-4.

24. Chin, C.H., Chen, S.H., Wu, H.H., Ho, C.W., Ko, M.T., and Lin, C.Y. (2014). cytoHubba: identifying hub objects and sub-networks from complex interactome. BMC Syst Biol 8 *Suppl 4*, S11. 10.1186/1752-0509-8-S4-S11.

25. Chinnery, P.F., Zwijnenburg, P.J., Walker, M., Howell, N., Taylor, R.W., Lightowlers, R.N., Bindoff, L., and Turnbull, D.M. (1999). Nonrandom tissue distribution of mutant mtDNA. Am J Med Genet 85, 498–501.

26. Inak, G., Rybak-Wolf, A., Lisowski, P., Pentimalli, T.M., Juttner, R., Glazar, P., Uppal, K., Bottani, E., Brunetti, D., Secker, C., et al. (2021). Defective metabolic programming impairs early neuronal morphogenesis in neural cultures and an organoid model of Leigh syndrome. Nat Commun 12, 1929. 10.1038/s41467-021-22117-z.

27. Qian, X., Su, Y., Adam, C.D., Deutschmann, A.U., Pather, S.R., Goldberg, E.M., Su, K., Li, S., Lu, L., Jacob, F., et al. (2020). Sliced Human Cortical Organoids for Modeling Distinct Cortical Layer Formation. Cell Stem Cell 26, 766–781 e769. 10.1016/j.stem.2020.02.002.

28. Scholle, L.M., Zierz, S., Mawrin, C., Wickenhauser, C., and Urban, D.L. (2020). Heteroplasmy and Copy Number in the Common m.3243A>G Mutation-A Post-Mortem Genotype-Phenotype Analysis. Genes (Basel) 11. 10.3390/genes11020212.

29. Miyahara, H., Tamai, C., Inoue, M., Sekiguchi, K., Tahara, D., Tahara, N., Takeda, K., Arafuka, S., Moriyoshi, H., Koizumi, R., et al. (2023). Neuropathological hallmarks in autopsied cases with mitochondrial diseases caused by the mitochondrial 3243A>G mutation. Brain Pathol 33, e13199. 10.1111/bpa.13199.

30. Iwata, R., Casimir, P., Erkol, E., Boubakar, L., Planque, M., Gallego Lopez, I.M., Ditkowska, M., Gaspariunaite, V., Beckers, S., Remans, D., et al. (2023). Mitochondria metabolism sets the species-specific tempo of neuronal development. Science 379, eabn4705. 10.1126/science.abn4705.

31. Iwata, R., Casimir, P., and Vanderhaeghen, P. (2020). Mitochondrial dynamics in postmitotic cells regulate neurogenesis. Science 369, 858–862. 10.1126/science.aba9760.

32. Karppa, M., Herva, R., Moslemi, A.R., Oldfors, A., Kakko, S., and Majamaa, K. (2005). Spectrum of myopathic findings in 50 patients with the 3243A>G mutation in mitochondrial DNA. Brain 128, 1861–1869. 10.1093/brain/awh515.

33. Dolle, C., Flones, I., Nido, G.S., Miletic, H., Osuagwu, N., Kristoffersen, S., Lilleng, P.K., Larsen, J.P., Tysnes, O.B., Haugarvoll, K., et al. (2016). Defective mitochondrial DNA homeostasis in the substantia nigra in Parkinson disease. Nat Commun 7, 13548. 10.1038/ncomms13548.

34. Baker, A., Kalmbach, B., Morishima, M., Kim, J., Juavinett, A., Li, N., and Dembrow, N. (2018). Specialized Subpopulations of Deep-Layer Pyramidal Neurons in the Neocortex: Bridging Cellular Properties to Functional Consequences. J Neurosci 38, 5441–5455. 10.1523/JNEUROSCI.0150-18.2018.

35. Hevner, R.F. (2007). Layer-specific markers as probes for neuron type identity in human neocortex and malformations of cortical development. J Neuropathol Exp Neurol 66, 101–109. 10.1097/nen.0b013e3180301c06.

36. Romito-DiGiacomo, R.R., Menegay, H., Cicero, S.A., and Herrup, K. (2007). Effects of Alzheimer’s disease on different cortical layers: the role of intrinsic differences in Abeta susceptibility. J Neurosci 27, 8496–8504. 10.1523/JNEUROSCI.1008-07.2007.

37. Braak, H., Del Tredici, K., Rub, U., de Vos, R.A., Jansen Steur, E.N., and Braak, E. (2003). Staging of brain pathology related to sporadic Parkinson’s disease. Neurobiol Aging 24, 197–211. 10.1016/s0197-4580(02)00065-9.

38. Braak, H., Rub, U., Schultz, C., and Del Tredici, K. (2006). Vulnerability of cortical neurons to Alzheimer’s and Parkinson’s diseases. J Alzheimers Dis 9, 35–44. 10.3233/jad-2006-9s305.

39. Pfisterer, U., Petukhov, V., Demharter, S., Meichsner, J., Thompson, J.J., Batiuk, M.Y., Asenjo-Martinez, A., Vasistha, N.A., Thakur, A., Mikkelsen, J., et al. (2020). Identification of epilepsy-associated neuronal subtypes and gene expression underlying epileptogenesis. Nat Commun 11, 5038. 10.1038/s41467-020-18752-7.

40. Leone, D.P., Srinivasan, K., Chen, B., Alcamo, E., and McConnell, S.K. (2008). The determination of projection neuron identity in the developing cerebral cortex. Curr Opin Neurobiol 18, 28–35. 10.1016/j.conb.2008.05.006.

41. Haast, R.A.M., Ivanov, D., RJT, I.J., Sallevelt, S., Jansen, J.F.A., Smeets, H.J.M., de Coo, I.F.M., Formisano, E., and Uludag, K. (2018). Anatomic & metabolic brain markers of the m.3243A>G mutation: A multi-parametric 7T MRI study. Neuroimage Clin 18, 231–244. 10.1016/j.nicl.2018.01.017.

42. Haast, R.A.M., De Coo, I.F.M., Ivanov, D., Khan, A.R., Jansen, J.F.A., Smeets, H.J.M., and Uludag, K. (2022). Neurodegenerative and functional signatures of the cerebellar cortex in m.3243A > G patients. Brain Commun 4, fcac024. 10.1093/braincomms/fcac024.

43. Szklarczyk, D., Nastou, K., Koutrouli, M., Kirsch, R., Mehryary, F., Hachilif, R., Hu, D., Peluso, M.E., Huang, Q., Fang, T., et al. (2025). The STRING database in 2025: protein networks with directionality of regulation. Nucleic Acids Res 53, D730–D737. 10.1093/nar/gkae1113.

