## Supplementary figures and images for "Mitochondrial DNA heteroplasmy drives cortical neuronal disturbances in human organoids harbouring the common m.3243A>G mutation"

### Figure S1

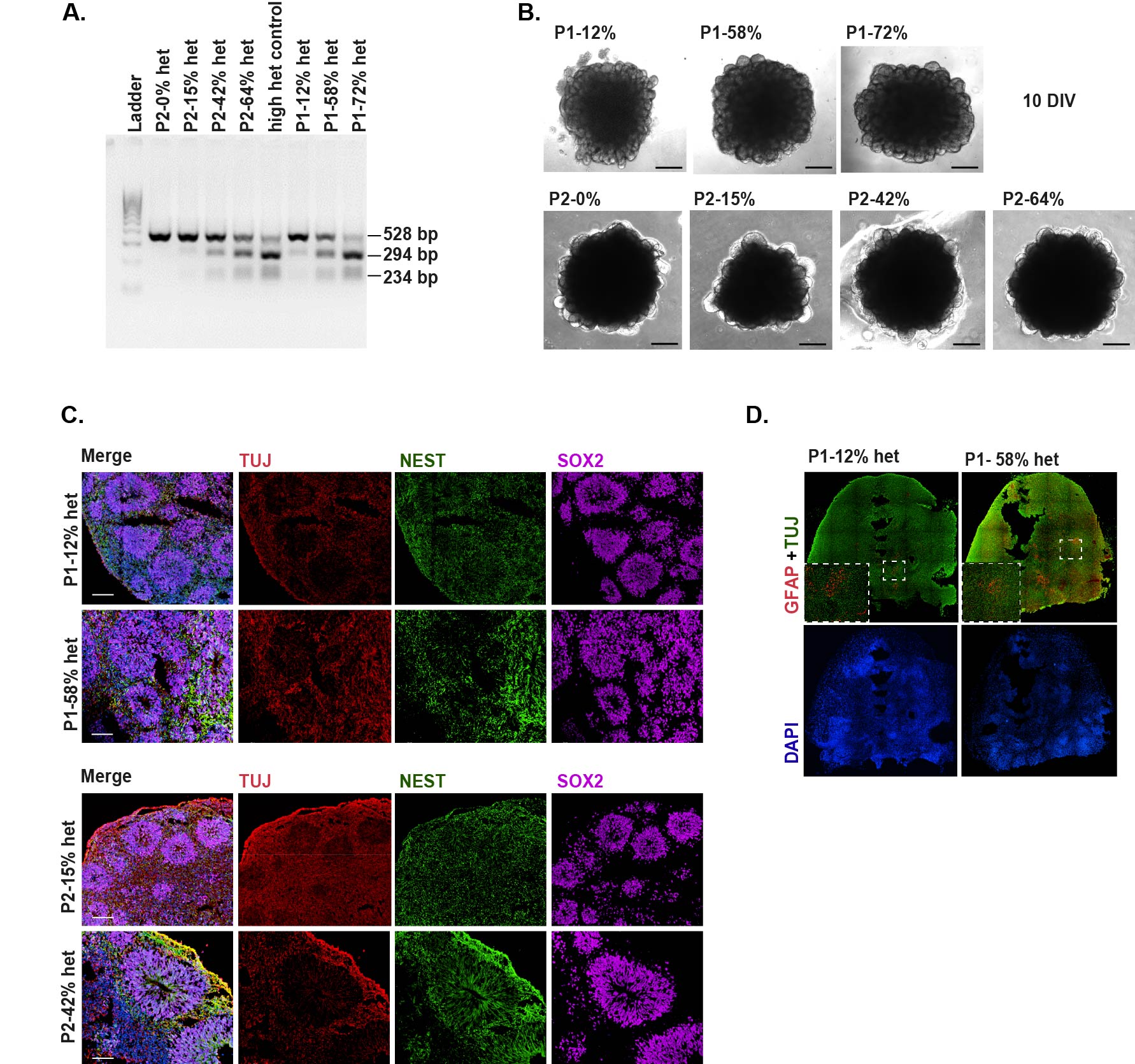

### Figure S2

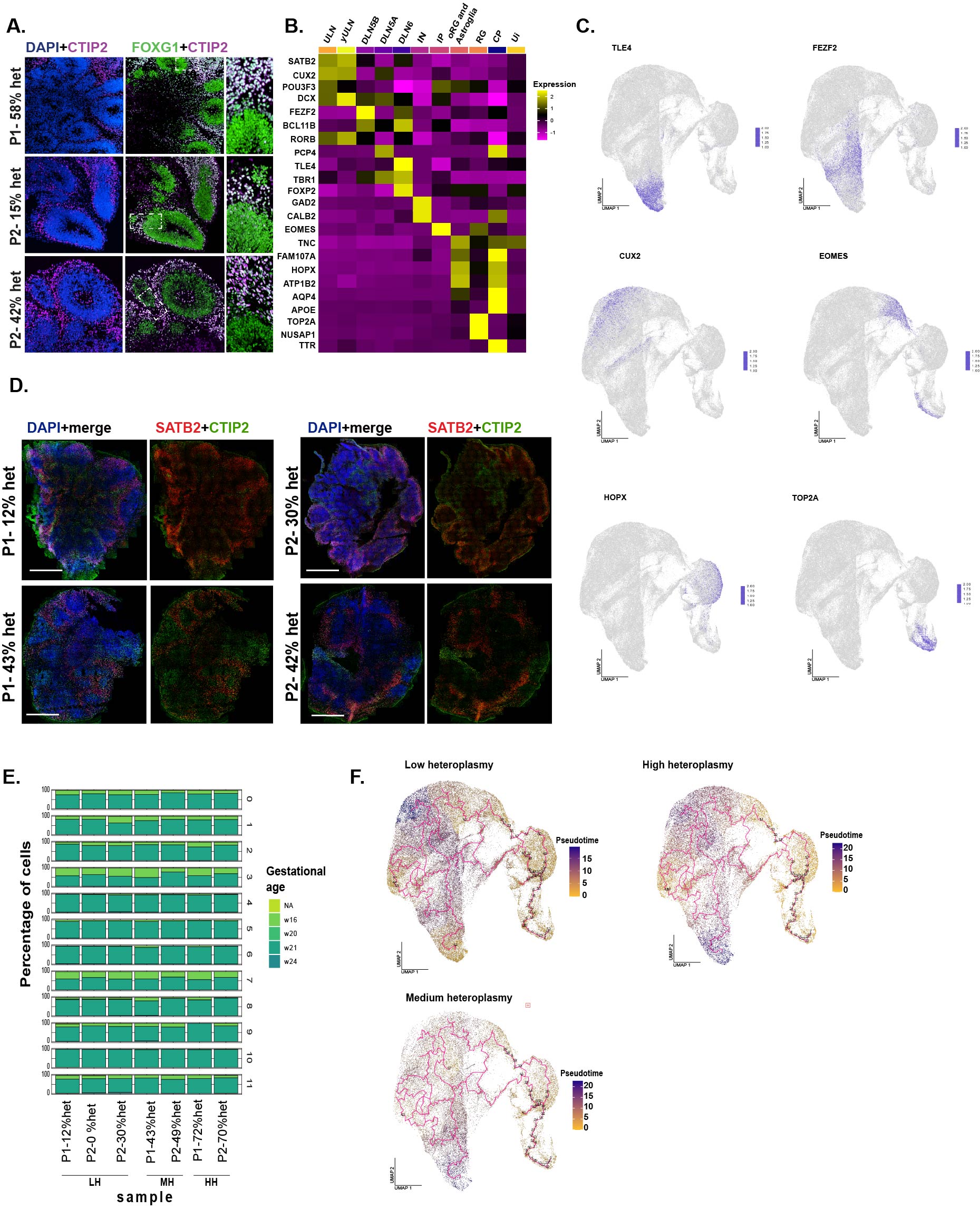

### Figure S3

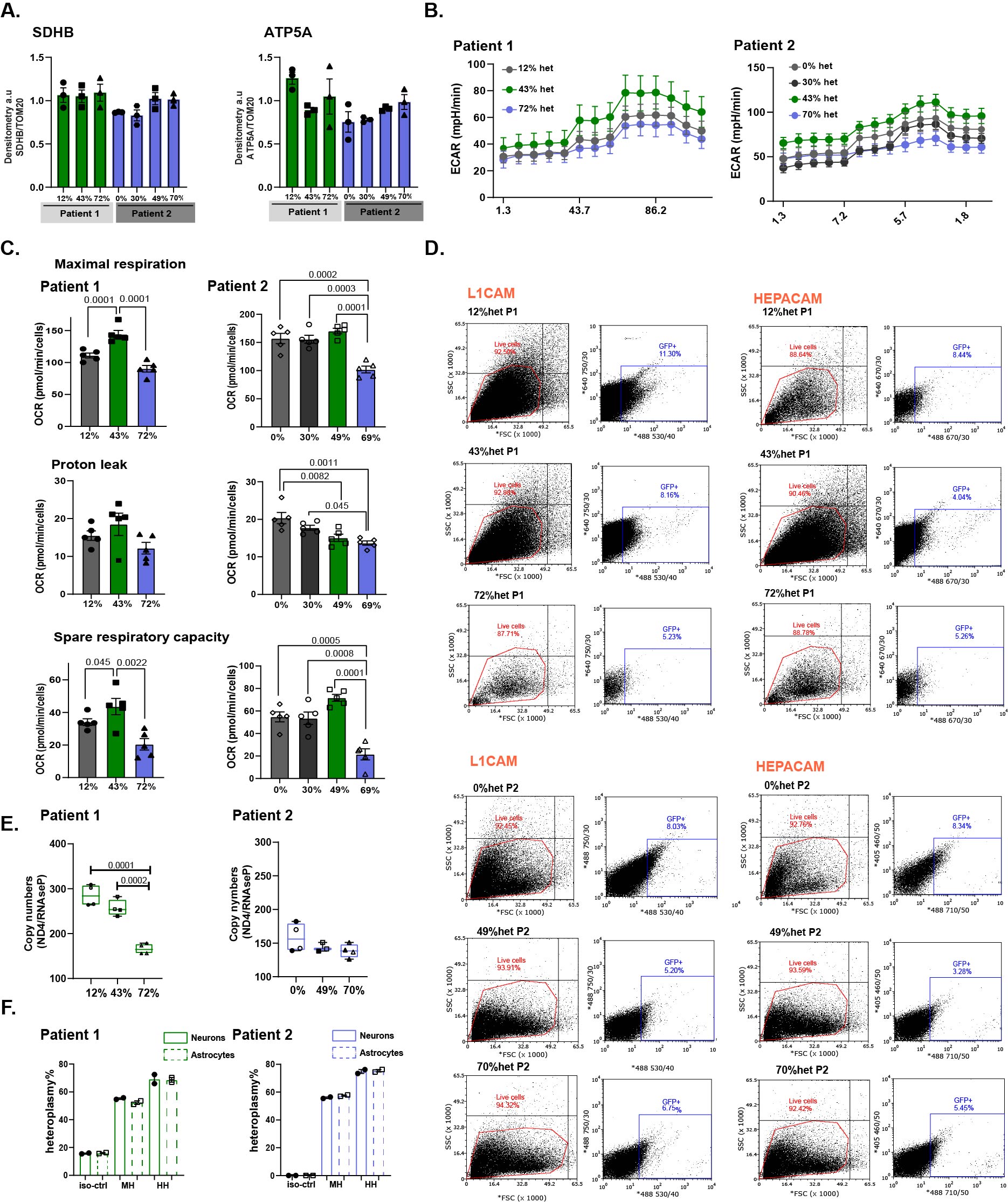

### Figure S4

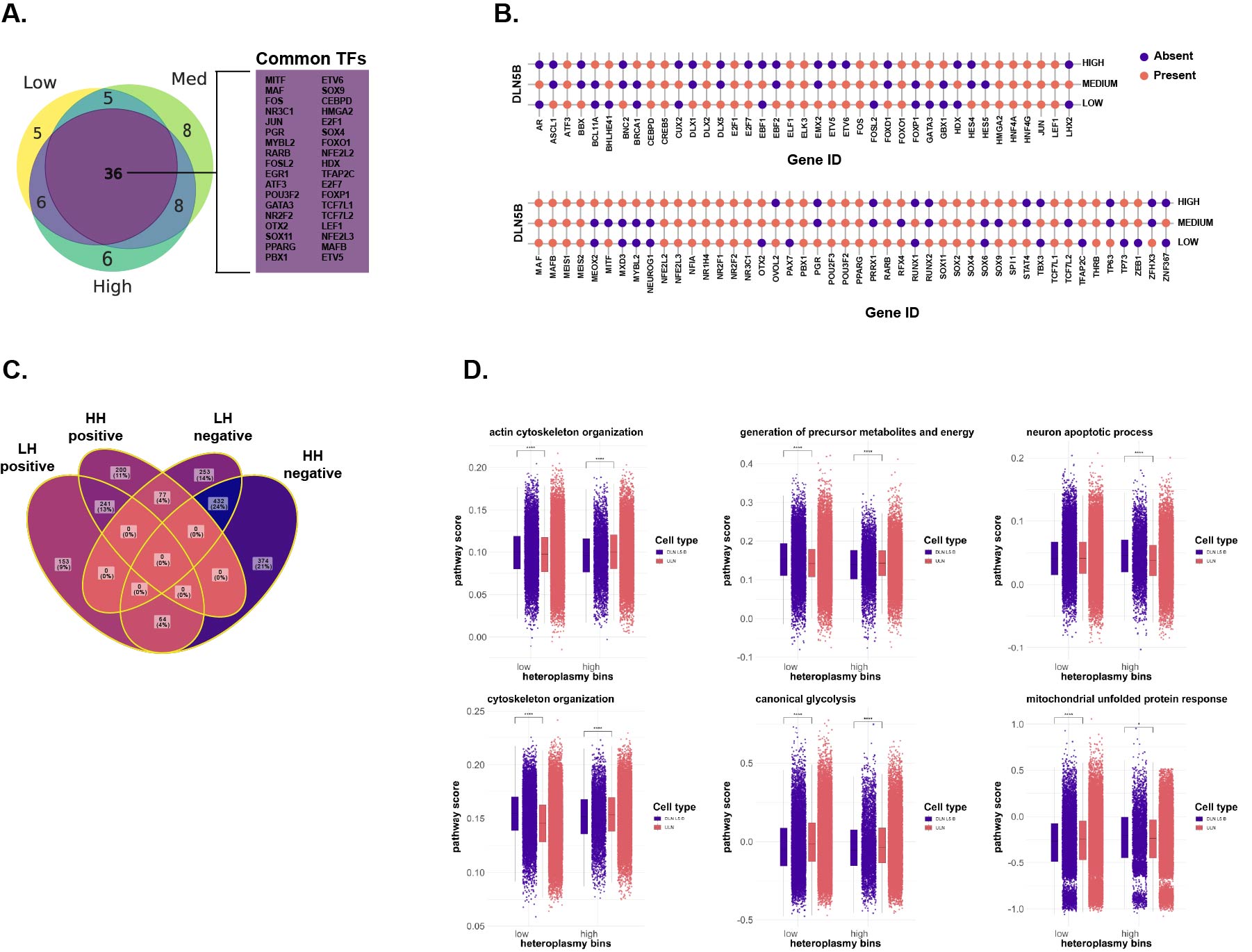
